# SARS-CoV2 infection triggers reactive astrocyte states and inflammatory conditions in long-term Human Cortical Organoids

**DOI:** 10.1101/2024.04.16.589036

**Authors:** M. Colinet, I. Chiver, A. Bonafina, G. Masset, D. Almansa, E. Di Valentin, J.C. Twizere, L. Nguyen, I. Espuny-Camacho

## Abstract

SARS-CoV2, severe acute respiratory syndrome coronavirus 2, is frequently associated with neurological manifestations. Despite the presence of mild to severe CNS-related symptoms in a cohort of patients, there is no consensus whether the virus can infect directly brain tissue or if the symptoms in patients are a consequence of peripheral infectivity of the virus. Here, we use long-term human stem cell-derived cortical organoids to assess SARS-CoV2 infectivity of brain cells and unravel the cell-type tropism and its downstream pathological effects. Our results show consistent and reproducible low levels of SARS-CoV2 infection of astrocytes, deep projection neurons, upper callosal neurons and inhibitory neurons in 6 months human cortical organoids. Interestingly, astrocytes showed the highest infection rate among all infected cell populations that led to increased presence of reactive states. Further, transcriptomic analysis revealed overall changes in expression of genes related to cell metabolism, astrocyte activation and, inflammation and further, upregulation of cell survival pathways. Thus, local and minor infectivity of SARS-CoV2 in the brain may induce widespread adverse effects and may lead to resilience of dysregulated neurons and astrocytes within an inflammatory environment.

## Introduction

SARS-CoV2, severe acute respiratory syndrome coronavirus 2, that causes coronavirus disease 2019 (COVID-19), is associated with a plethora of symptoms in patients ranging from mild fever to pneumonia and, in a small proportion of cases, systemic defects that can lead to the death of the individual^1–4^. Recent findings have associated SARS-CoV2-infectivity related effects with alterations in multiple organs including lung, heart, kidney, and brain. While SARS-CoV2 mainly targets the respiratory system, neurological manifestations have also been detected in patients. These neurological adverse effects include anosmia, strokes, seizures and impaired consciousness, among others^5–8^. Despite the observation of brain related symptoms in a fraction of the patients, the analysis of postmortem brains or cerebral spinal fluid (CSF) samples from surviving patients has led to contradictory results with most reports failing to detect the presence of SARS-CoV2 particles^9–13^. These observations suggest either low levels of infection in brain tissue or on the contrary an absence of infectivity in the brain, where neurological-related effects would rely on SARS-CoV2 peripheral organ infection.

SARS-CoV2 viral entry into target cells has been proposed to be mediated by the biding of the spike S protein to the receptor angiotensin converting enzyme 2 (ACE2). This event is followed by the activation of the spike S protein by a cleavage enacted by the host proprotein convertase furin and by membrane-associated serine proteinases (MASPs)^14,15^, which leads to the fusion of the virus membrane with the host membrane^2,16^. Whereas high levels of ACE2 expression have been detected in major target tissues of SARS-CoV2 infection such as lung, kidney and heart, ACE2 levels are reported to be low or undetectable in the brain^17^. However, the expression of ACE2 co-receptors that potentiate SARS-CoV2 infectivity has been reported in a number of tissues including the brain. These co-receptors include neuropilin 1 (NRP1) cell surface receptor^16,18–20^, dypeptidylpeptidase 4 (DPP4)^21^, tyrosine-protein kinase receptor UFO (AXL)^22^, the transmembrane glycoprotein (CD147/BSG)^7,23,24^, stress-inducible ER chaperone (GRP78/HSPA5)^25,26^. These reports support the possibility that alternative receptors and co-receptors are mediating SARS-CoV2 entry in brain cells.

Concerning the downstream effects of SARS-CoV2 infectivity, recent reports performing multi-organ screenings by magnetic resonance imaging (MRI) have detected multiple anomalies in lungs and brain in a cohort of COVID-19 patients 6 months post-infection^27^. In addition, postmortem analysis of patient brain tissue has revealed broad cellular perturbations of most cell types by single-cell transcriptomic^28^ and proteomic^9^ analysis. These studies suggested the preferential infectivity of astrocyte glial-cells and their consequent change towards a higher reactive state linked to higher levels of inflammation detected in the brain. Whereas most studies have been focused on adult brain, a recent report has shown hemorrhagic phenotypes in the cortex of human fetuses exposed to SARS-CoV2^29^, through a mechanism related to reduction in blood vessel integrity^29–31^.

Several studies have used human pluripotent stem cell-derived brain organoids to analyze the infectivity of SARS-CoV2 in brain and its pathological downstream effects. Infectivity of brain cell types has been reported to be low compared to the levels of infection detected for instance in models including choroid plexus epithelial and ependymal cells^32,33^ and perivascular pericyte-like cells^34^. A few studies have corroborated *in vitro* the infectivity of astrocytes by SARS-CoV2 in agreement with the reports from postmortem brain sample analysis^19,35^. However, systematic studies analyzing the percentage of cells with different identities infected by the virus and their susceptibility for SARS-CoV2 infection in human cortical organoid models at different stages *in vitro* are currently missing.

Here, we analyze neural cell infectivity of SARS-CoV2 in human cortical organoid models to understand the cellular tropism of the virus at different brain organoid maturation stages. We found that astrocytes, deep projection neurons, upper callosal neurons and inhibitory neurons were infected by SARS-CoV2. Astrocytes showed the highest rate of infected cells among the specific cell population in human cortical organoids at late stages *in vitro*. We detected low levels of ACE2 expression but higher levels of some of its co-receptors, including NRP1, CD147, GRP78, NRP2 and AXL, which may be involved in viral entry in the target cells. Mechanistically, we found that upon SARS-CoV2 infection, astrocytes acquire reactive states leading to broad adverse effects on cortical organoid cell populations by triggering an inflammatory response counteracted by upregulation of cell survival pathways.

## Results

### SARS-CoV2 virus consistently infects human cortical organoids albeit at low levels

We aimed to test the infectivity and to identify the downstream effects of SARS-CoV2 viral infection in the human brain by means of an *in vitro* human cortical organoid model (hCO)^36,37^. For this purpose, hCOs^37^ of 2, 3.5 and 6 months *in vitro* (2M, 3.5M, 6M) were incubated for 3h with SARS-CoV2 viruses^38^ at a 0.5 multiplicity of infection (MOI) previously reported for assessing SARS-Cov2 infectivity of brain organoids^32^ and analyzed the results 1 day (24h) or 3 days (72h) after infection (Fig. 1A). Immunolabelings of hCOs revealed the presence of SARS-CoV2 nucleocapside (NC) proteins in less than 0.1% of hCO cells (Fig. 1B-K and Movie S1). No differences were observed between the different hCO stages analyzed for the number of cells infected (NC+), nor between different times after infection (Fig. 1J). We reasoned that the exposition of hCOs to a higher viral load would result in increased cell infectivity. Our results showed that doubling the MOI of SARS-CoV2 exposure led to a higher level of infectivity of hCOs, suggesting that SARS-CoV2 infection follows a dose-dependent response (Fig. 1L). In contrast, and as expected, a similar approach using kidney Vero cells, a susceptible cell type for SARS-CoV2 infection^39^, resulted in a higher level of infectivity at all SARS-CoV2 multiplicities of infection tested (Fig. S1). Next, we analyzed the level of expression of known SARS-CoV2 receptors and co-receptors in the hCOs. RNA sequencing of late stage (6M) hCOs confirmed low expression of the major SARS-CoV2 receptor, ACE2^2,16,17^, and higher expression of some of the SARS-CoV2 co-receptors such as NRP1, CD147/BSG; GRP78/HSPA5; DPP4 and AXL^7,16,18–26^ as well as some of the enzymes involved in the cleavage of the SARS-CoV2 virus spike protein to mediate viral entry to the cell, such as transmembrane serine protease 2 (TMPRSS2), cathepsin B (CTSB), cathepsin L (CTSL) and furin^14,23^ (Fig. 1M). Immunolabelings confirmed the low/absence of expression of ACE2 in mouse cortex samples and in brain organoids (Fig. S2A). In contrast, ACE2 expression was detectable in kidney Vero cells^39^ (Fig. S2B). In agreement with our RNAseq data we could detect the expression of NRP1 in brain organoids and in mouse cortex (Fig. S2C). These results indicate that human cortical organoids are susceptible to infection by the SARS-CoV2 virus albeit at low levels, possibly due to the low levels of ACE2 present in the brain. Moreover, our results also suggest that some ACE2 co-receptors and proteases involved in entry of the SARS-Cov2 virus in the cells are expressed in human cortical organoids.

**Figure 1:**
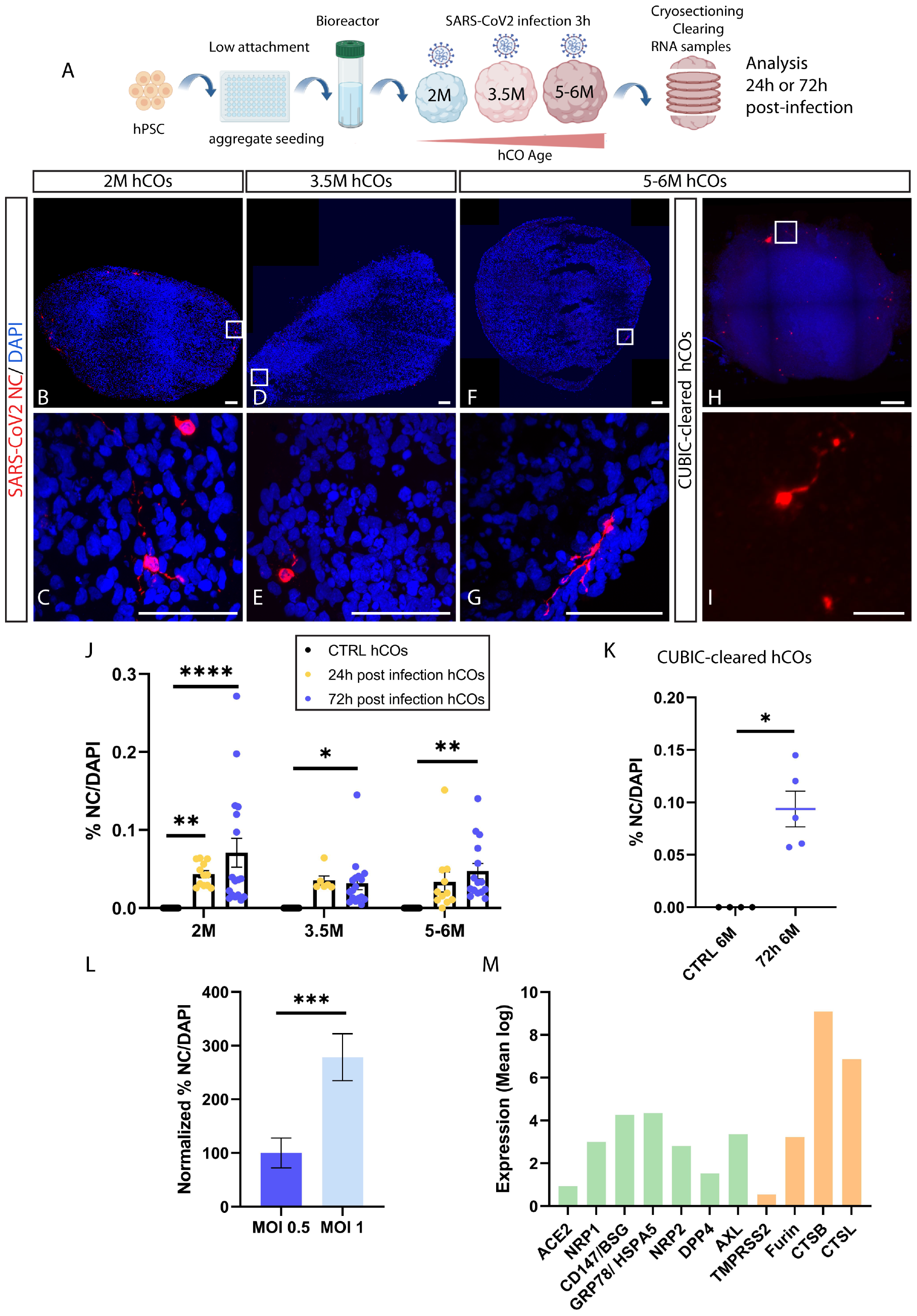
SARS-CoV2 infects human cortical organoids at different stages *in vitro*. (A) Schematic illustration of the procedures followed for derivation of human cortical organoids of different stages *in vitro* (2M, 3.5M and 5-6M), hCO infection by SARS-CoV2 viral particles, and subsequent analysis. Created with BioRender.com. (B-G) Tile-Scan (B,D,F) and single (C,E,G) confocal images of cryosections immunostained for the SARS-CoV2 antigen nucleocapside (NC, in red) in 72h SARS-Cov2 post infection hCOs at all time points (2M, 3.5M, 5-6M). (H-I) Tile-Scan (H) and single (I) light-sheet microscope images of clarified (CUBIC) 72h SARS-CoV2 post infected hCOs and immunostained for NC (in red) at 6M. Counterstaining was performed using DAPI. (J) Quantification of the percentage of NC+ cells among the total number of DAPI cells, in control (CTRL), 24h and 72h SARS-CoV2 post-infection hCOs at 2M, 3.5M and 5-6M *in vitro*. Data are represented as mean percentages ± SEM (3-4 differentiations: CTRL n=15-18; 24h n=6-12; 72h n=15-17). Two way ANOVA with Tukey’s multiple comparison test *p<0.05 **p<0.01 ****p<0.0001. (K) Quantification of the percentage of NC+ cells 72h post infection in whole clarified (CUBIC) 72h SARS-CoV2 6M hCOs. Data are represented as mean percentages ± SEM (CTRL n=4; 72h n=5). Mann-Whitney test *p<0.05. (L) Quantification of the percentage of NC+ cells in hCOs exposed to SARS-CoV2 MOI 0.5 and MOI 1 at 2M. Data are represented as mean percentages ± SEM (MOI 0.5 n=6; MOI 1 n=3). Mann-Whitney test ***p<0.001. (M) Expression levels (mean log) of SARS-CoV2 receptors and co-receptors (in green), and of SARS-CoV2 viral entry-related proteases (in orange) in 6M hCOs (n=4) from bulk RNAseq. Data are represented as mean ± SEM. Scale bars: 100µm (B,D,F); 50µm (C,E,G); 300µm (H) and 40 µm (I).

### Deep layer projection neurons and astrocytes are the major cell types infected by SARS-CoV2 virus in hCOs at late stages *in vitro*

Next, we sought to analyze the cell identity of SARS-CoV2 infected cells in hCOs at different stages of maturation. First, we tested the infectivity of progenitors in hCOs from 2M up to 5-6M *in vitro*. Our experiments showed negligible numbers of either PAX6 or KI67 progenitor/proliferating cells infected by SARS-CoV2 in hCOs at any of the time points tested (Fig. S3A-D). Contrarily, we found that significant numbers of CTIP2+ deep layer projection neurons, and to a lesser extent CUX1+ upper layer cortical callosal neurons were infected in hCOs at early (2M, 3.5M), and later (5-6M) time points (Fig. 2A-D). Strikingly, hCOs at later stages also showed a significant number of calbindin+ (CALB+) positive interneurons^40^ and GFAP+ astrocytes infected (Fig. 2E-H). The overall analysis of the cell type identity among SARS-CoV2 infected cells in hCOs at a later time point revealed that ∼40% of the infected cells corresponded to GFAP+ astrocytes, followed by ∼25% CTIP2+ cells, ∼15% CALB+ cells and about ∼3% CUX1+ cells (Fig. 2I). Further, we looked into the percentage of cells infected among the total specific cell type population in hCOs at 6 months. This analysis revealed that astrocytes were the cell type showing the highest percentage of cells infected among the total number of the specific cell population, when compared to excitatory and inhibitory neurons of the cortex (Fig. 2J).

**Figure 2:**
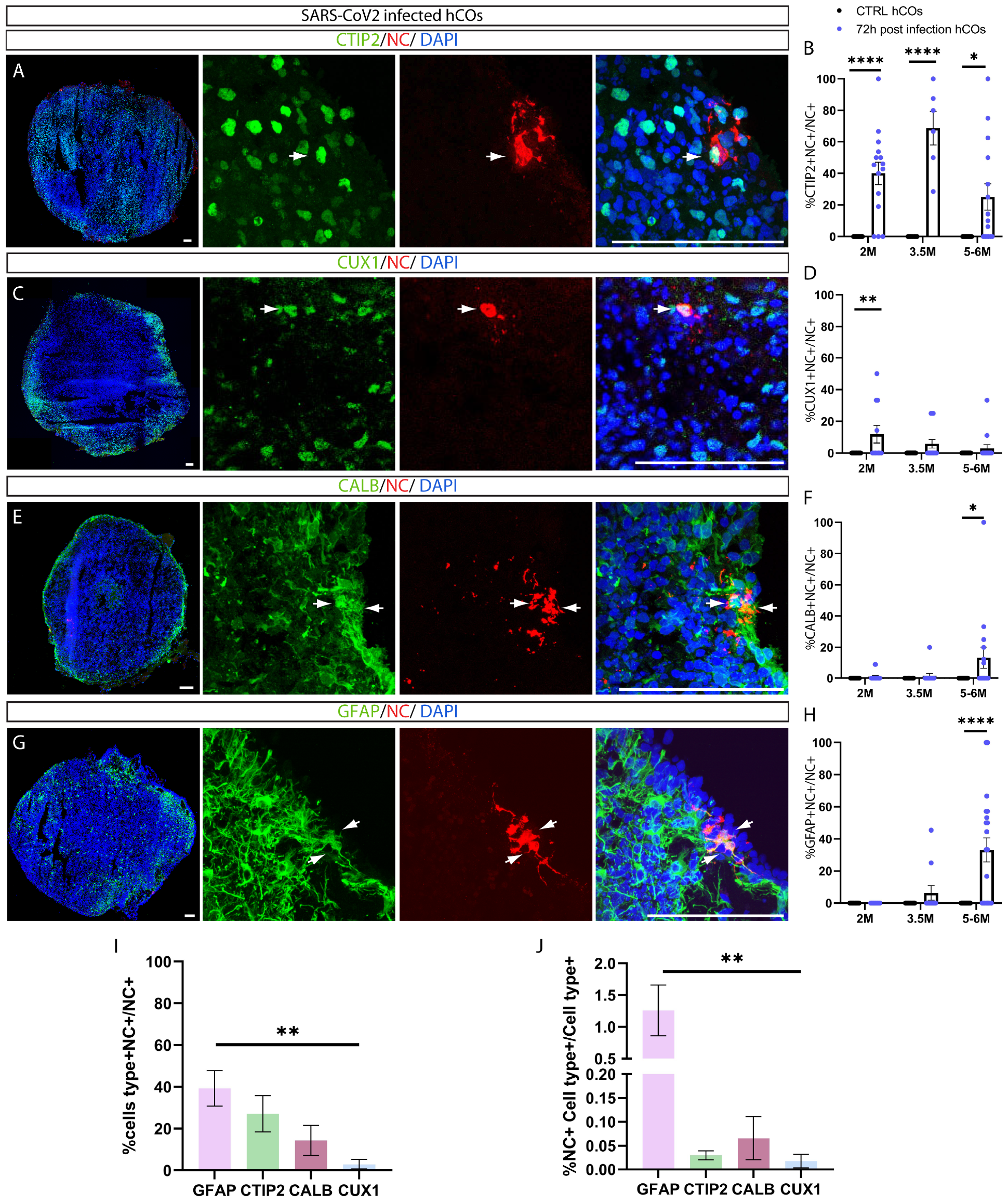
SARS-CoV2 infects mostly astrocyte cells at late stages in hCOs. (A,C,E,G) Tile-Scan (left panel) and single confocal images of cryosections immunostained for the SARS-CoV2 antigen nucleocapside (NC, in red) and the deep layer cortical neuronal marker CTIP2 (in green, A); the upper layer cortical neuronal marker CUX1 (in green, C); the interneuron marker Calbindin (in green, E), or the astrocyte marker GFAP (in green, G) in 6M 72h post-infection hCOs, and high magnification insets. Counterstaining was performed using DAPI. (B,D,F,H) Quantification of the percentage of double positive CTIP2+NC+ (B); CUX1+NC+ (D); CALB+NC+ (F); GFAP+NC+ (H), among the total NC+ population in 2M, 3.5M and 5-6M 72h SARS-CoV2 hCOs compared to control (CTRL). Data are represented as mean percentages ± SEM (2-3 differentiations: CTRL n=8-20 72h n=11-20. Two way ANOVA with Tukey’s multiple comparison test *p<0.05 **p<0.01 ****p<0.0001. (I) Percentage of specific cell types double positive for NC among the total NC+ population in 5-6M 72h SARS-CoV2 hCOs. Data are represented as mean percentages ± SEM (3-4 differentiations: 72h n=14-16). Two way ANOVA with Tukey’s multiple comparison test **p<0.01. (J) Percentage of NC+ infected cell type among the total numbers of the cell type-specific population in 5-6M 72h SARS-CoV2 hCOs. Data are represented as mean percentages ± SEM (3-4 differentiations: 72h n=15-20. Two way ANOVA with Tukey’s multiple comparison test *p<0.05 **p<0.01. Scale bar: 100µm (A,C,E,G). White arrows show cells double positive for NC and the specific cell-type marker tested (A,C,E,G).

These results suggest that SARS-CoV2 can infect astrocytes, deep projection neurons, upper callosal neurons and inhibitory neurons in the brain and identify astrocytes as the cell type with the highest susceptibility for infection in hCOs.

### hCO cells infected with SARS-CoV2 do not undergo cell death

We asked whether SARS-CoV2 infectivity would lead to a higher rate of cell death in hCOs. To answer this question we performed immunolabeling to detect activated cleaved-caspase 3 (CASP3) after 24 hours or 72 hours post exposure to SARS-CoV2, at different hCOs maturation stages in culture. Our analysis revealed no difference in the number of cleaved-CASP3 positive cells between infected and control hCOs (Fig. 3A-C). Moreover, we found no cleaved-CASP3 positive cells infected by the virus, further suggesting that the few apoptotic events detected were not SARS-CoV2-dependent (Fig. 3D). On the contrary, as shown before, Vero cells showed high levels of cleaved-CASP3 following SARS-CoV2 exposure^41^ (Fig. S4). We next sought to examine cell density as a global measurement of cell death in hCOs, as shown previously^42^. Our analysis showed no change in cell density following SARS-Cov2 infection (Fig. 3E-F).

**Figure 3:**
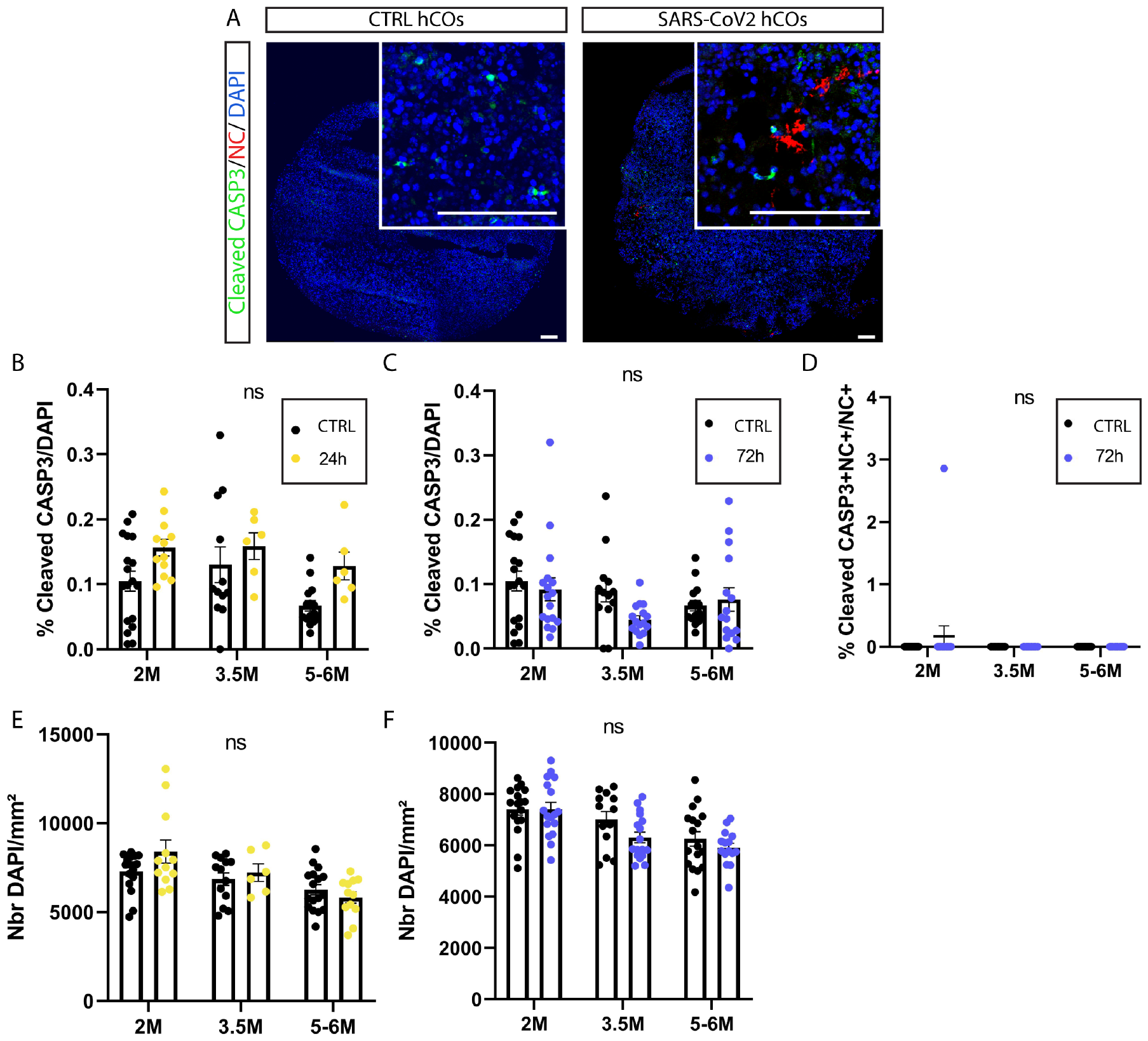
SARS-CoV2 infectivity does not induce cell death in hCOs. (A) Tile-Scan confocal images of cryosections immunostained for the apoptotic marker caspase 3 (CASP3)+ cells (in green) and the SARS-CoV2 antigen NC (in red) in 5-6M CTR and 72h hCOs anf high magnification pics (insets). Counterstaining was done with DAPI. (B-C) Quantification of the percentage of CASP3+ cells among the total number of cells DAPI+ in CTRL and 24h post infection hCOs (B) or in CTRL and 72h post infection hCOs (C). Data are represented as mean percentages ± SEM (3 differentiations: 2M n= 12-18; 3.5M n=6-16; 5-6M n=6-15). Two-way ANOVA with Tukey’s multiple comparison test, ns= non-significant. (D) Quantification of the percentage of double positive cells for CASP3 and NC among the NC+ population at all stages. Data are represented as mean percentages ± SEM (3 differentiations: 2M n= 17-18; 3.5M n= 13-16; 5-6M n= 15). Two-way ANOVA with Tukey’s multiple comparison test, ns= non-significant. (E-F) Quantification of the cell density (number of DAPI cells per mm^2^) in CTRL, 24h and 72h post infection hCOs at all stages. Data are represented as mean percentages ± SEM (3 differentiations: 2M n=12-17; 3.5M n=6-17; 5-6M n=12-17). Two-way ANOVA test with Sidak multiple comparison test. ns: non-significant. Scale bar: 100µm (A-B).

As we detected a preferential cell-type tropism for SARS-CoV2 virus combined with low levels of infectivity, we reasoned that certain hCO cell populations could be more affected than others, and/or that effects on cell death could be present in a localized manner specifically around SARS-CoV2 infected areas in hCOs. To unravel cell type-specific effects inside hCOs following SARS-CoV2 infection, we measured the percentage of different cell types following SARS-CoV2 infection and compared them to control. We found no change in the percentage of proliferative progenitors (KI67+), deep and upper cortical layer neurons (CTIP2+, CUX1+) or interneurons (CALB+)^40^ in hCOs at 72h post infection compared to control (Fig. S3E-H). Next, we measured cell density and nuclei area within SARS-CoV2 infected regions (NC+) in hCOs to unravel localized effects. Neither nuclei density, nor nuclei area inside NC+ areas were affected following SARS-CoV2 infection (Fig. S5A-C), suggesting no cell-type or region-specific-dependent effects on cell death in infected hCOs.

These results suggest that infection by SARS-CoV2 does not trigger cell apoptosis in hCOs at different maturation stages in culture.

### SARS-CoV2 virus infects different brain regional identities *in vitro*

In order to test the tropism of SARS-CoV2 in the central nervous system, we infected human brain organoids with rostral and caudal regional identities. For this purpose, we generated brain organoids (BOs) with ventral telencephalic medial ganglionic eminences (MGE) identity, which are enriched in cortical interneurons and their progenitors^43–45^; BOs that model the thalamus (THL)^46^; and some that recapitulate features of the cerebellum (CRB)^47^. We exposed hCOs, MGE BOs, THL BOs and CRB BOs to SARS-CoV2 virus MOI 0.5 and measured its effects after 72h post infection *in vitro*. Our analysis revealed that MGE BOs had the highest percentage of NC+ cells compared to hCOs, THL BOs or CRB BOs (Fig. 4A-G). Next, we measured the level of apoptosis-related events in those different brain regional identity BOs by measuring the percentage of activated-CASP3+ cells. This analysis revealed that MGE BOs, THL BOs and CRB BOs showed no differences in the percentage of cleaved-caspase3 cells compared to their controls, similarly to the previous data obtained with hCOs (Fig. 4H).

**Figure 4:**
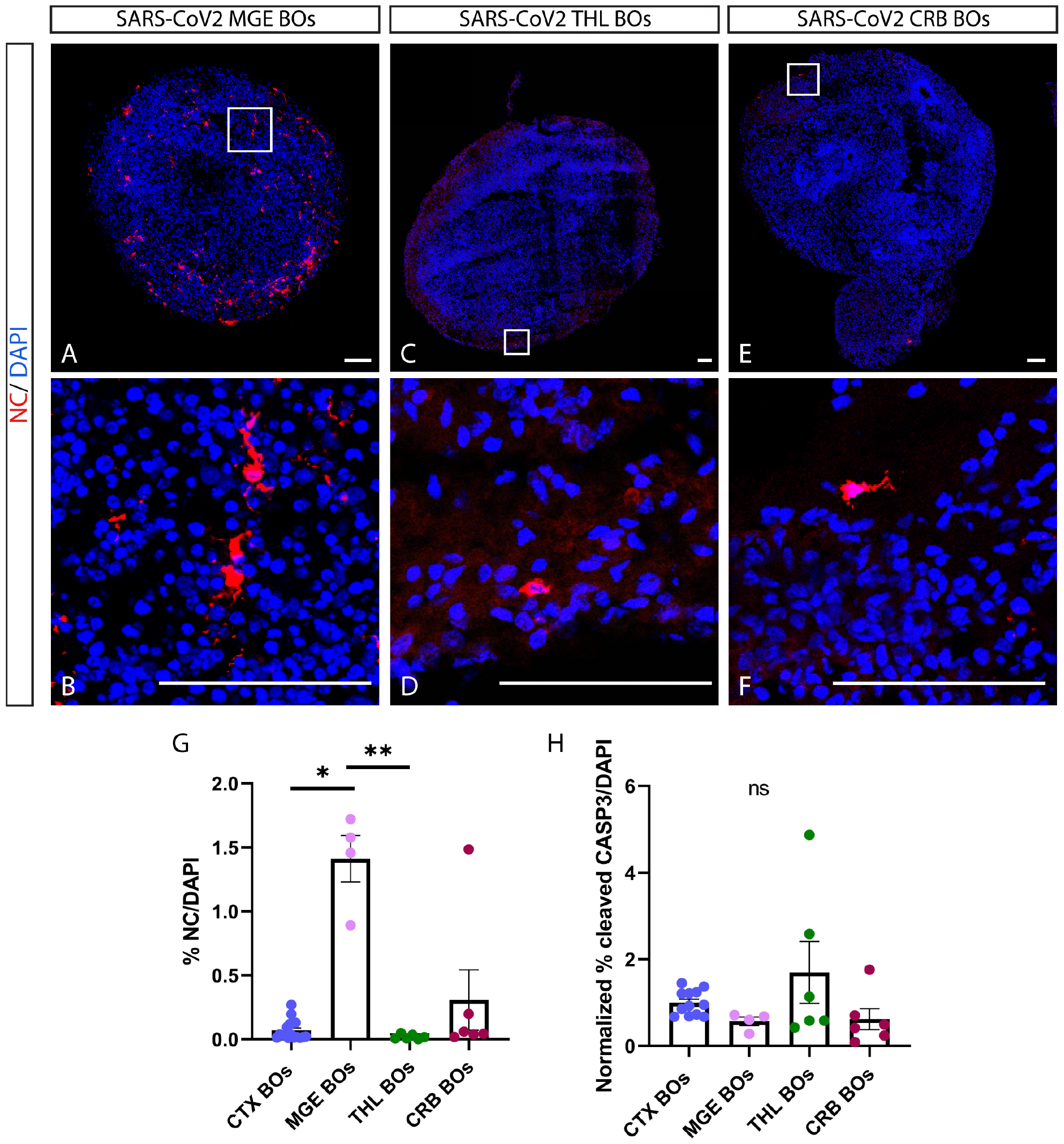
SARS-CoV2 virus infects different brain regional identity BOs in vitro. (A-F) Tile-Scan (A,C,E) and single (B,D,F) confocal images of cryosections immunostained for the SARS-CoV2 antigen nucleocapside (NC, in red) in 72h MGE BOs (A-B), THL BOs (C-D) and CRB BOs (E-F) at 2M. Counterstaining was performed using DAPI. (G) Quantification of the percentage of NC+ cells among the total number of cells DAPI+ in CTX hCOs, MGE BOs, CRB BOs and THL BOs. Data are represented as mean percentages ± SEM (CTX n=17; MGE n= 4; CRB n= 6; THL n=6). Kruskal-Wallis test with Dunn’s multiple comparison test.. *p < 0.05; **p < 0.01. (H) Quantification of the relative percentage of CASP3+ cells among the total number of cells DAPI+ in SARS-CoV2 infected CTX hCOs, MGE BOs, CRB BOs and THL BOs compared to CTRL CTX hCOs, MGE BOs, CRB BOs and THL BOs expressed as 1. Data are represented as normalized mean normalized percentage ± SEM (CTX n=17; MGE n= 4; CRB n= 6; THL n=6). Kruskal-Wallis test with Dunn’s multiple comparison test. Ns= non-significant. Scale bar: 100µm (A-F).

These results suggest that rostral forebrain telencephalic, diencephalic, and caudal hindbrain regions of the brain show susceptibility to SARS-CoV2 infection; with MGE identities showing the highest number of cells infected *in vitro*.

### Infection of hCOs by SARS-CoV2 virus results in phenotypic changes in the astrocyte population that correlate with higher reactive states

Since astrocytes are the most frequent cell type infected in hCOs, we hypothesized that their infectivity by SARS-CoV2 would trigger phenotypic and functional changes. We first analyzed the impact of SARS-CoV2 infection in the percentage of astrocyte (GFAP+) coverage in infected hCOs as compared to control. This analysis revealed that infected hCOs had a lower GFAP+ organoid area coverage as compared to controls (Fig. 5A-D, G). While this phenotype did not result from the reduction in number of astrocytes in infected hCOs (Fig. 5H), we noticed that infected astrocytes were smaller. Reactive astrocytes have indeed been shown to change their morphology and size according to their reactive state *in vitro* and *in vivo*^48,49^. Accordingly, our analysis revealed that in SARS-CoV2 infected astrocytes in hCOs had a reduced area as compared to control astrocytes (Fig. 5E-F, I), suggesting a higher astrocyte reactive state upon infection.

**Figure 5.**
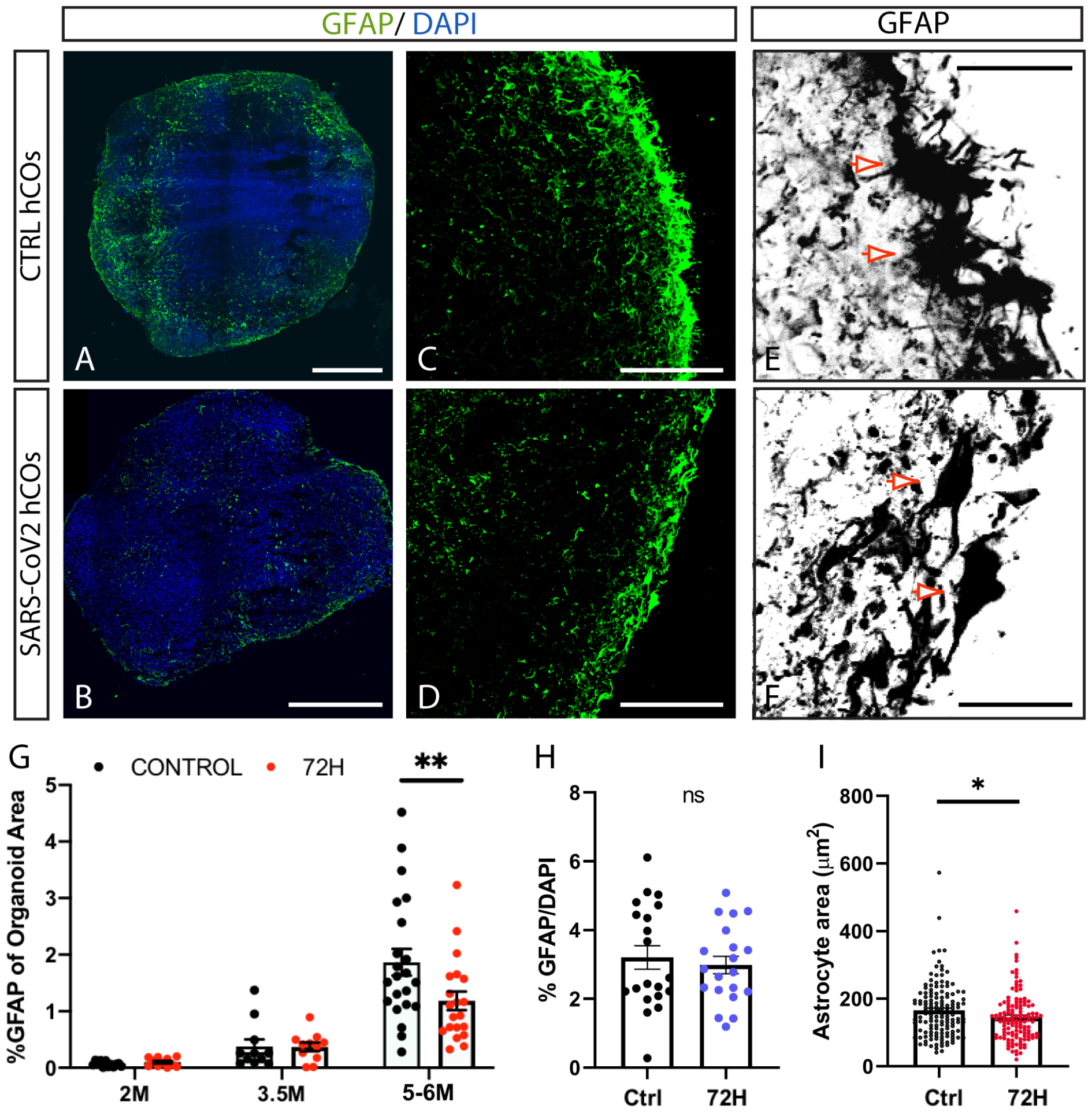
Astrocytes show increased reactive states following SARS-CoV2 infection. (A-F) Tile-Scan (A-B) and single (C-F) confocal images of cryosections immunostained for the astrocyte marker GFAP (in green: A-D; black and white mask: E-F) in 5-6M CTRL and 72h hCOs. Counterstaining was performed using DAPI. (G) Percentage of organoid area covered by GFAP+ cells in CTRL and 72h hCOs of 2M, 3.5M and 5-6M. Data are represented as mean percentages ± SEM (2-4 differentiations: 2M n= 6-12; 3.5M n= 6-11; 5-6M n= 12-22). Two-Way ANOVA Sidak’s multiple comparison test **p<0.01. (H) Quantification of the percentage of GFAP+ cells among the total number of cells DAPI+ in CTRL and 72h 5-6M hCOs. Data are represented as mean percentages ± SEM (4 differentiations: CTRL n= 20; 72h n=20). Unpaired t-test ns=non-significant. (I) Quantification of astrocyte area in CTRL and 72h 5-6M hCOs. Data are represented as mean percentages ± SEM. (4 differentiations: CTRL n= 139 astrocytes from 23 organoids; 72h n=128 from 20 organoids). Mann-Whitney two-tailed test * p<0.05. Scale bar 500µm (A-B); 100 µm (C-D); 25µm (E-F). Red arrows show examples of representative astrocytes (E,F).

Altogether, these results suggest that infection of astrocytes by SARS-CoV2 triggers their reactive state in hCOs.

### SARS-CoV2 infection in hCOs increases the expression of pro-inflammatory and pro-survival genes

We hypothesized that higher reactive state in astrocytes triggered by SARS-CoV2 infection could have overall effects in cell populations in hCOs. To test this hypothesis, we analyzed 5-6M infected and control hCOs by bulk RNA sequencing to unravel genes and pathways affected following infection of SARS-CoV2. First, we validated the expression of SARS-CoV2 viral genes in infected hCOs (Fig. 6A). We did not detect statistical changes in the expression of SARS-CoV2 receptor and co-receptors following SARS-CoV2 infection of hCOs (Fig. S6A). Next, we analyzed genes differentially expressed between infected and control hCOs (padj <0.05). We found that several genes related to astrocyte-mediated inflammation such as alpha 1 antichymotrypsin (SERPINA3)^50,51^, S100 calcium binding protein A10 (S100A10)^52,53^ and the interleukin 13 receptor subunit alpha1 (IL13RA1), the immune system-related gene complement factor I (CFI), and the epidermal growth factor receptor (EGFR) were dysregulated following SARS-CoV2 infection in hCOs. In addition, the expression of lipid-metabolism related genes, which are associated to astrocyte reactive states^48^, such as phospholipase C delta 3 (PLCD3) and phospholipase C delta 1 (PLCD1) were also affected in infected hCOs. Interestingly, instead of effects promoting cell death pathways we detected an upregulation of the cell survival-related genes superoxide dismutase 2 (SOD2)^54^, brain and acute leukemia cytoplasmic protein (BAALC)^55,56^, the CD44 astrocyte receptor^57–59^ and the cytokine hepatocyte growth factor (HGF)^60^ following SARS-CoV2 infection (Fig. 6B-C and Table S1). Top candidate pathways (enrichR) and gene set enrichment analysis (GSEA) revealed the link of oxidative phosphorylation, glycolysis, TNFA signaling via NFKB, protein secretion, inflammatory responses, epithelial-mesenchymal transition, and lipid metabolism-related pathways (Fig. 6D-E, Fig. S6D and Tables S2 and S3). Interestingly, and in agreement with our data, the expression of genes related to glycolysis and lipid metabolism, as well as the pathway epithelial-mesenchymal transition, have been shown to be modulated upon a change towards a more reactive state in astrocytes^48,49^.

**Figure 6:**
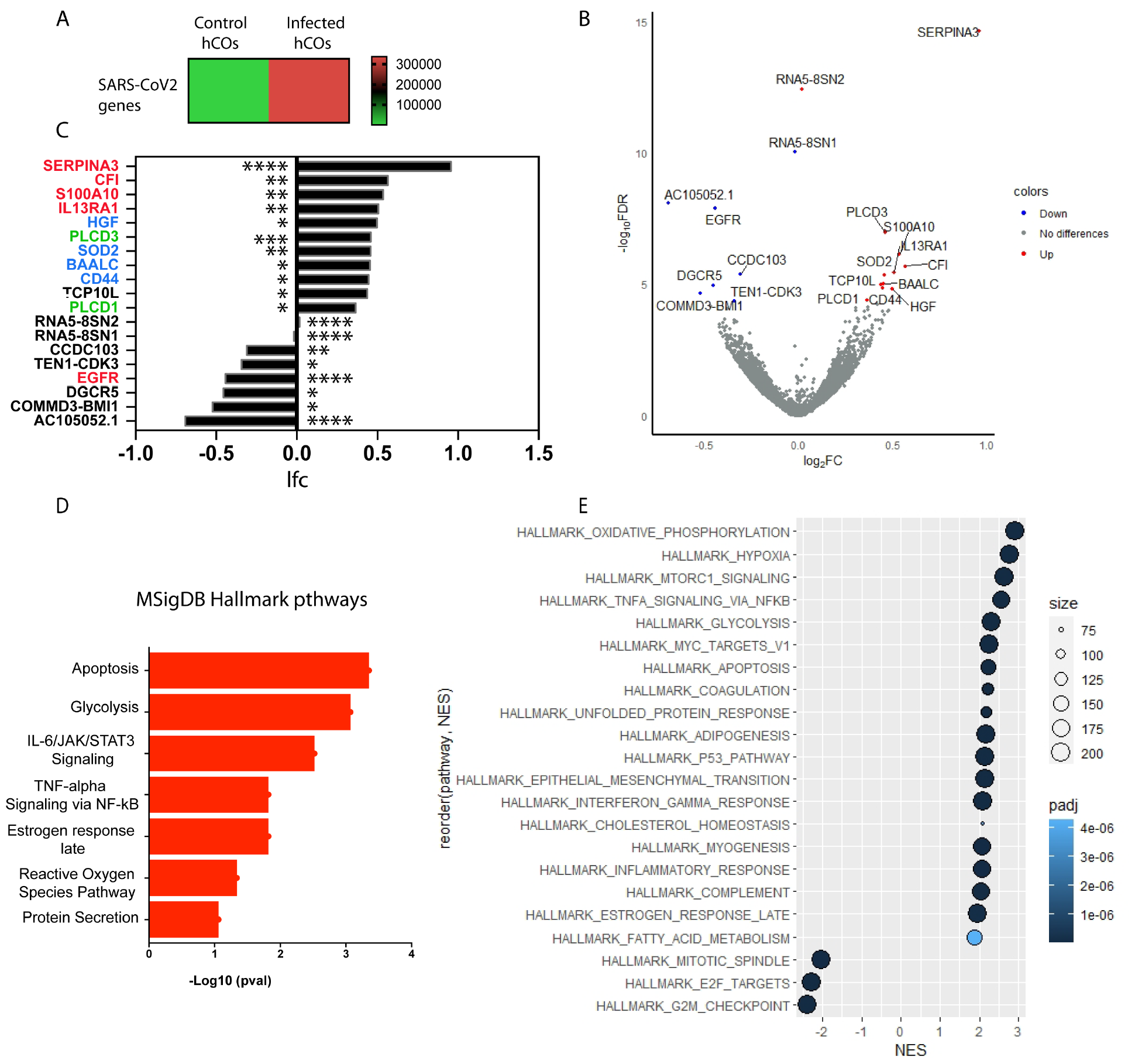
SARS-CoV2 infection of 5-6M cortical organoids triggers immune system and oxidative stress related responses. (A) Heatmap representation of SARS-CoV2 normalized counts in CTRL or 72h SARS-CoV2 5-6M hCOs. (B) Volcano plot representation of the changes in genes expression (DEG) in 5-6M 72h SARS-CoV2 post infection hCOs compared to CTRL hCOs, representing the Log2foldChange (log_2_FC) and the p_adj_ value (-log_10_FDR). Genes with a p_adj_ value below 0.05 and with a negative Log2foldChange are represented in blue (downregulated genes), and genes with a p_adj_ value below 0.05 with a positive Log2foldChange are represented in red (upregulated). Genes that have a p_adj_ value above 0.05 are represented in grey (non significantly changed). (C) Bar graph representation of the log fold change (lfc) of the differentially expressed genes in 72h hCOs compared to CTRL hCOs. In red: Astrocyte-mediated inflammatory genes; in green: lipid metabolism-related genes; in blue: cell survival-related genes. (D) Representation of the top pathways from MSigDB Hallmark pathways database based on the significance of enrichment of upregulated and downregulated DEGs. (E) Bubble plot of the Gene Set Enrichment Analysis (GSEA) representing the enriched pathways with a p_adj_ value < 0.0001. Pathways that have a normalized enrichment score (NES) positive >0 are upregulated whereas pathways with a NES negative <0 are downregulated. Pathways were ordered by the NES score in a decreasing manner. Size represented the number of genes from the RNAseq presented in the pathway. All data were generated from bulk RNAseq analysis of 2 independent differentiations (CTRL n=4; 72h n=4). *p < 0.05; **p < 0.01; ***p < 0.001; ****p < 0.0001.

We further analyzed the expression of candidate pro-inflammatory serpin family A member 3 (SERPINA3), and pro-survival SOD2 genes in infected hCOs, as they could have important implications in the inflammatory cascades mediated by reactive astrocytes following SARS-CoV2 infection in hCOs. Our analysis revealed a higher number of cells expressing the reactive astrocyte marker SERPINA3 and the pro-survival gene SOD2 in hCOs 72h post infection compared to control (Fig. 7A-B, E-F). Moreover, we specifically found a higher percentage of astrocytes expressing SOD2 in hCOs 72h post infection compared to control astrocytes, suggesting that astrocytes in infected hCOs switch on the expression of the SOD2 gene as a mechanism against viral infection to promote anti-apoptotic pathways (Fig. 7C, G). In line with this observation, we found a lower percentage of cells labeled for gamma H2AX, a marker of DNA damage in the cell (Fig. 7D, H).

**Figure 7.**
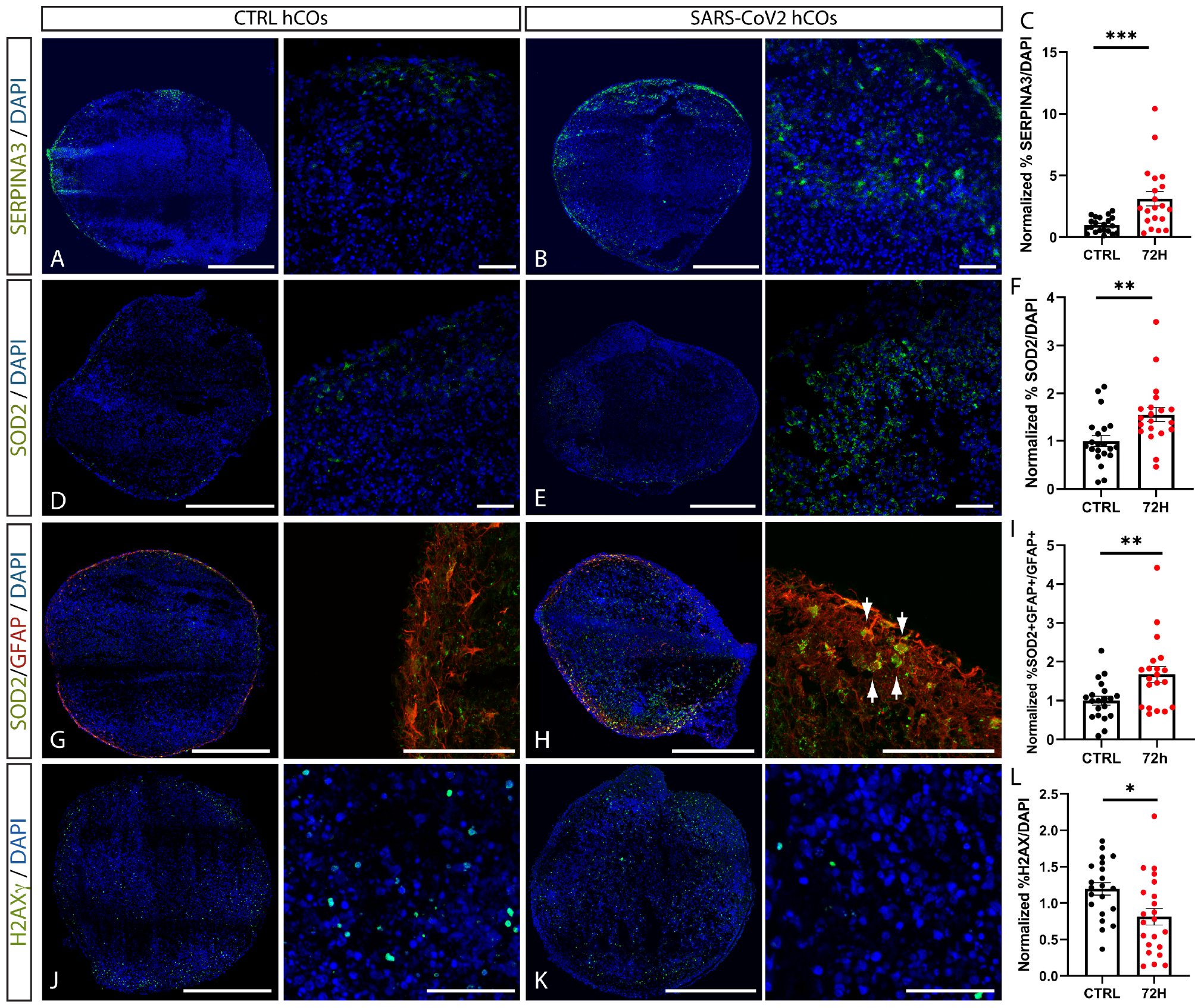
Increased expression of astrocyte reactivity markers and pro-survival genes in infected hCOs. Tile-Scan (A,D,G,J) and single (B,E,H,K) confocal images of cryosections immunostained for the reactive astrocyte marker SERPINA3 (in green; A-B), the pro-survival gene SOD2 (in green; D-E,G-H), the astrocyte marker GFAP (in red; G-H), and the DNA double strand break marker gamma H2AX (in green; J-K) in 5-6M CTRL and 72h hCOs. Counterstaining was performed using DAPI. White arrows show cells double positive for SOD2 and GFAP (H). (C,F,I,L) Quantifications of the percentage of SERPINA3+ (C), SOD2+ (F), SOD2+GFAP+ (I) and gamma H2AX+ (L) cells over the total number of DAPI+ cells in control and 72h 5-6M hCOs. Data are normalized by the mean percentage area value of each trial. Data are represented as mean percentages ± SEM (4 differentiations: CTRL n= 20; 72h n=20). (C,F) Mann-Whitney test **p<0.01, ***p<0.001 (I) Mann-Whitney test **p<0.01. (L) Two-tailed t-test *p<0.05. Scale bars: 500µm (A-K) and 50 µm (higher magnification views on the right).

These results suggest that SARS-CoV2 infection triggers a higher reactive state in astrocytes leading to an overall increased expression of pro-inflammatory genes that could precede an upregulation of pro-survival pathways to prevent cell death mechanisms in hCOs.

## Discussion

The understanding of the ability of the virus SARS-CoV2 to infect the human brain has so far remained elusive due to the fact that only a few studies have reported the presence of viral RNA in CSF from COVID19 patients or detected the presence of the virus in postmortem brain samples^9,30,61,62^. However, neurological manifestations are present in about at least a third of COVID19 patients suggesting the possibility that pathological effects are triggered in the brain of patients^63^. Lack of SARS-CoV2 particle detection in the brain may result from the challenge to access CSF and/or brain tissue from COVID19 patients during critical phases of SARS-CoV2 infection, possibly combined with a low level of infectivity of SARS-CoV2 in the brain when compared to other tissues such as the respiratory system. In support of this hypothesis, a recent study based on 11 autopsies from COVID19 patients at different stages of the disease and using novel technologies for RNA detection (droplet digital PCR) succeeded in showing the presence of SARS-CoV2 RNA in 10 out of 11 of the brain samples analysed^10^. However, a limitation of this and similar previous studies is that patient cohorts may have been more susceptible for SARS-CoV2 infectivity and its pathological downstream effects than control groups, due to preexistent unrelated medical conditions that finally led to their death from severe COVID-19. Moreover, previous concerns are the contamination of brain samples with blood cells or lung tissue during the autopsy procedure that could lead to false positive detection of SARS-CoV2. The aim of the current study is to analyze the ability of SARS-CoV2 to infect the brain, to distinguish its tropism towards different brain cell types, and to unravel major downstream effects of SARS-CoV2 infection in brain using a long-term human *in vitro* cortical organoid model.

The infectivity of SARS-CoV2 has been previously assessed in human telencephalic organoids derived from default differentiation paradigms and compared to that in choroid plexus organoids. This study found mostly infectivity of SARS-CoV2 restricted to choroid plexus cells with absence or minimum detection of virus RNA or protein in other brain organoid cell types^32^. Later reports using different brain organoid models reported however infectivity of the virus in brain cells besides choroid plexus cells^33,64–68^. Most of these studies reported a similar infectivity rate among cell types, including progenitors, neurons and glia cells, or on the contrary, higher infectivity in neurons^33,64–66,68,69^. Some recent studies have suggested a specific cellular tropism of SARS-CoV2 for astrocytes in the brain^19,35^. However a systematic and quantitative analysis of brain cell types susceptible to SARS-CoV2 infection in brain organoids and the analysis of different brain organoid maturation stages *in vitro* including long-term cultures has not yet been performed. Here we present an exhaustive analysis of regionalized brain organoids at 3 different *in vitro* stages up to 6 months in culture exposed to SARS-CoV2 virus that reveals consistent infectivity of the virus at all stages albeit low levels (around 0.1%). This low percentage of infectivity in cortical organoids was consistent with the low level of detection of ACE2 at the mRNA level. Nevertheless, we could detect higher expression of some of the ACE2 co-receptors such as NRP1 and DPP4 and some of the proteases involved in viral entry to the cell. In contrast with previous reports we have not found infectivity of progenitors or proliferative cells in hCOs at any of the maturation stages analyzed. Our analysis revealed a time-dependent pattern of tropism of the virus where at early stages deep projection neurons and upper callosal neurons were the most infected cell types whereas at late time points there was a switch towards astrocytes as the main target of the SARS-CoV2 virus. While the percentage of SARS-CoV2 infected cells represented around 0.1% of all hCO cells, SARS-CoV2 infected more than a 1% of the total astrocyte population at late stages *in vitro*, and this percentage was far higher than the one reported for any other infected cell type, suggesting a more specific tropism of SARS-CoV2 for astrocytes.

In this study we have compared the efficiency of SARS-CoV2 to infect brain organoid models that mimic different regions of the forebrain and the developing cerebellum. Intriguingly, we found that ventralized organoids (MGE) were more susceptible to SARS-CoV2 infection as compared to other regionalized organoids. Other reports have previously interrogated the infectivity of SARS-CoV2 using multiple brain organoid regionalities *in vitro*, however, these studies did not reveal any difference in the percentage of infectivity between organoid models^33^. Our findings suggest that cortical interneurons and/or their progenitors could be important targets of SARS-CoV2 infection. Further work is needed to assess whether cortical interneurons that are traveling towards the cortex are also susceptible to SARS-CoV2 infection and whether this would affect their migration in forebrain assembloids^43–45^.

The study of the downstream consequences of SARS-CoV2 infection in the brain has been the focus of several works for instance those reporting MRI scans-based brain abnormalities such as lesions in the white matter in COVID19 patients 6 months post-infection^27^. However, the extent of those brain anatomical alterations did not correlate with the neurological symptoms of the patients and there was an age mismatch between the control and infected group that could lead to the anatomical differences detected. Conversely, recent studies have found no major cytopathological effects outside the respiratory tract in postmortem patient samples^10^. Previous *in vitro* and *in vivo* studies have shown an increased cell death rate among SARS-CoV2 infected cells in the brain^9,19,35,70^. However, these results are in contrast with other reports showing no major cell loss in different models^69,71^. We also aimed to clarify SARS-CoV2 downstream effects using cortical brain organoids at different stages *in vitro*. Our in depth analysis showed no major effects in cell death: no changes in apoptotic cell number, no changes in overall cell density, nor in specific cell-type numbers, nor effects on nuclei size in SARS-Cov2 infected areas or within neighboring regions in infected cortical organoids. While no major change of cell survival was detected upon SARS-CoV2 infection of hCOs, we observed cellular changes in the astrocyte population suggestive of a higher reactive state. Moreover, RNAseq performed on infected hCOs revealed an overall increased expression of inflammatory genes such as SERPINA3, increased oxidative phosphorylation, TNFA signaling, astrocyte activation and metabolic changes, which suggests broad inflammatory-induced conditions in hCOs. Surprisingly, instead of cell death mechanism activation we detected upregulation of cell survival pathways (SOD2-related), which suggest that compensatory pro-survival mechanisms are triggered in astrocytes and neuronal populations following SARS-CoV2 infection of hCOs. In agreement with this, we detected lower levels of DNA damage marked by H2AX gamma in hCOs infected by the virus, indicating possibly more efficient repair mechanisms for double strand DNA break. These results could imply the appearance of resilient dysregulated astrocytes and neurons within an inflammatory context following SARS-CoV2 infection.

Our current study was limited to 2 short-range time points after SARS-CoV2 exposure, including 1 or 3 days, to understand the immediate major cellular and transcriptomic changes occurring in brain cell types as a direct cause of the infection of the virus. Recent studies have attempted to decipher the pathological events in COVID19 patients suffering from SARS-CoV2-related symptoms beyond 3 months after exposure to the virus, which is known today as long-COVID^72^. It would be thus interesting in future studies to assess long-term effects in brain cell types following SARS-CoV2 infection in cortical organoid models.

Lastly, some reports have hypothesized that SARS-CoV2-mediated effects in brain mimic the inflammatory conditions identified in the brains of patients suffering from neurodegenerative conditions^19,28,71^. Despite the limitation of our hCO model which lack major immune cell types, our results go in line with this hypothesis with pro-inflammatory changes triggered in our system only 1 to 3 days after infection of the virus combined with the activation of pro-survival pathways which could lead to survival of affected neurons and astrocytes. Strikingly, a known pro-inflammatory protein expressed in astrocytes, SERPINA3, which has been detected in plasma of middle-aged adults with high risk for dementia^73^, was also upregulated in our infected cortical organoids. One could envision cortical brain organoids infected with SARS-CoV2 as an interesting model to study susceptibility to develop dementia-like features after long-term culture that could parallel the *in vivo* situation of neurodegenerative patients.

In conclusion, we show here that hCOs at 3 different maturation stages *in vitro* up to 6 months present a reproducible level of low infectivity following SARS-CoV2 exposure. We demonstrate that astrocytes are the most representative cell type infected, that ultimately leads to a higher astrocyte reactive state and pro-inflammatory and cell survival-related changes in the overall brain cell type populations in cortical organoids. These data unravels important pathways triggered in the brain following SARS-CoV2 infection that could be relevant for the design of future therapies aiming at protecting the brain from inflammatory dementia-like related pathological changes.

## Conclusion

SARS-CoV2 infection leads to neurological symptoms in a subset of patients. Our study shows that SARS-CoV2 infects astrocytes, deep layer projection neurons, upper callosal neurons and interneurons of the cortex and trigger astrocyte reactive states and overall inflammation and cell survival pathways.

These findings could imply the emergence of resilient dysregulated neurons and astrocytes in the context of an inflammatory environment. Our model could be instrumental for the design of future strategies aiming at lowering SARS-CoV2-related neurological effects.

## Acknowledgements

This work was supported by the Fonds Nationale de la Recherche Scientifique (FNRS) Belgium: Credit Urgents de Recherche CUR 40002797 (to L. Nguyen, collaborator Espuny-Camacho), and PER 40003579 (to L. Nguyen, I. Espuny-Camacho, E. Di Valentin, J-C. Twizere). We thank the GIGA lentiviral Vectors, GIGA Imaging, and GIGA Genomics and Bioinformatics platforms for their contribution, help and support to this work.

## Author contributions

I.E-C designed the experimental plan. J-C.T and E.dV amplified the SARS-CoV2 virus and performed the SARS-CoV2 infections. M.C, I.E-C, I.C performed the experiments with the help of A.B, G.M and D.A. I.E-C, M.C, I.C analyzed all the results, I.E-C wrote the manuscript with the help of L.N, M.C, I.C and the rest of coauthors. I.E-C and L.N designed the initial study.

The authors declare no competing interests.

## Materials and Methods

### Human COs differentiation

Cortical brain organoids were developed following a modified version of the Sasai protocol^37^. hESC-H9 cell line (WA09) (Metadata: Female, no disease associated) was acquired from Wicell. H9 cells were dissociated using Accutase (Stem Cell Technologies: 7922) and 9000 cells were seeded in 96 well U bottom plates (Nunclon sphere: 15396123) per well. Cells were cultured in DMEM-F12 with 20% KO serum (ThermoFisher), supplemented with NEAA; penicillin/streptavidin and 100µm β-mercaptoethanol plus morphogens: 5µM SB (SB431542; Sigma) and 3µM IWR1 (Sigma) for 14 days. Rock inhibitor (Y-27632 2HC; Bio-Connect) was added until day 6 (20µM d0-d2; 10µM d2-d6). Organoids were cultured in DMEM-F12 medium supplemented with N2 (ThermoFisher: 17502048) and B27 (ThermoFisher: 11500446; 11530536) from day 14 and grown on orbital shakers (75 rpm) from day 21 and in bioreactor tubes (Bioké) from approximately day 46, speed 75 rpm. At day 70, 1% of matrigel (Corning: 11523550) was added to the culture. Medium was changed every 2 or 3 days.

### SARS-CoV2 infection

All experiments have been conducted by the GIGA viral platform (www.gigaviralvectors.uliege.be) in a Biosafety Level 3 lab using the SARS-CoV2 strain (BetaCov/Belgium/GHB-03021/2020 (EPI ISL 407976|2020-02-03)^38^ wild or recombinant SARS-CoV-2 strain harboring a mNeon-Green reporter^74^. VeroE6 cells were used to generate the SARS-CoV2 viral stock. Plaque assay method was used to determine the virus titer or plaque-forming unit (PFU)^32^.

### RNA sequencing, Data Analysis and Differential Expression Analysis

High quality RNA samples RIN values (8-10) were used for sequencing. Four samples of CTRL hCOs from 2 independent differentiations, and 4 samples from SARS-CoV2 72h post-infected hCOs from 2 different experiments were analyzed. The extracted RNA-seq libraries were generated by the GIGA Genomic platform (www.gigagenomics.uliege.be) using Illumina Stranded Total RNA Prep Ligation with Ribo-Zero Plus following the manufacturer protocol (Illumina ref: 20040529) and sequenced using the Illumina Novaseq 6000 sequencer, generating pair end 2×150bp reads. Downstream analysis was performed on R using DESeq2 method for normalization, clustering and differential expression analysis.

### Statistical analysis

All statistical analysis was performed using GraphPad (Prism). All data were first tested for normality with a Shapiro-Wilcoxon test and based on the result of this test, the parametric or corresponding non-parametric version of the test was used. The test for multiple comparisons was chosen based on the recommendation from GraphPad Prism. All information concerning the test used and the multiple comparisons test applied are found in the legend of the corresponding figure. All data presented are from at least 3 independent brain organoid batches.

