## Supplemental Materials for "SARS-CoV2 infection triggers reactive astrocyte states and inflammatory conditions in long-term Human Cortical Organoids"

**Supplemental Figures**

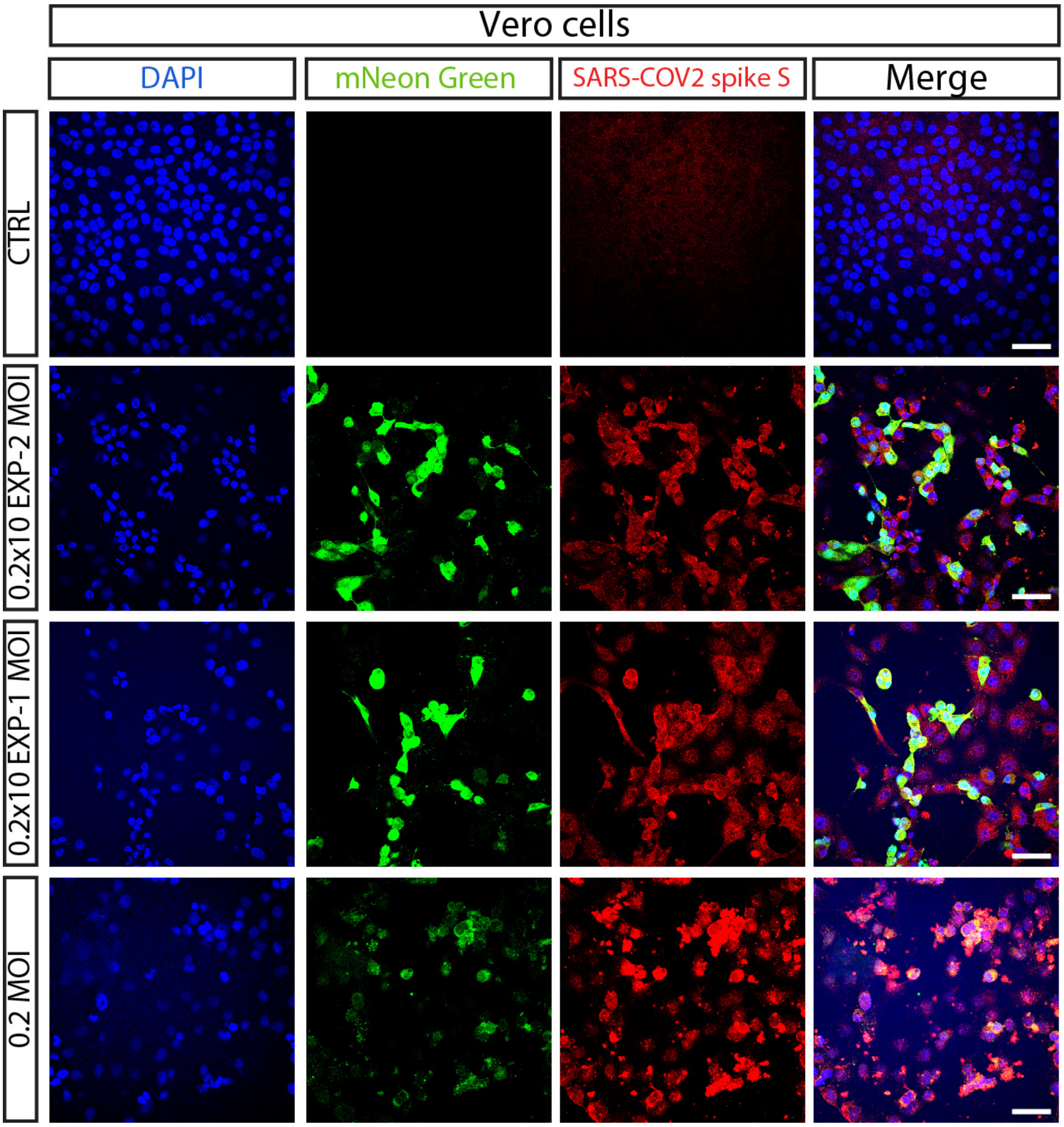


**Supplementary Figure S1. Vero cells show a high rate of infectivity following SARS-CoV2 virus exposure.**

Confocal images of CTRL and infected Vero cells with SARS-CoV2-mNeon Green modified virus (MOI: $0.5x10^{-2}$; $0.5x10^{-1}$; 0.5), immunostained with the SARS-CoV2 antigen spike S (in red) and mNeon Green (in green). Counterstaining was performed using DAPI. Scale bars: 50 μm

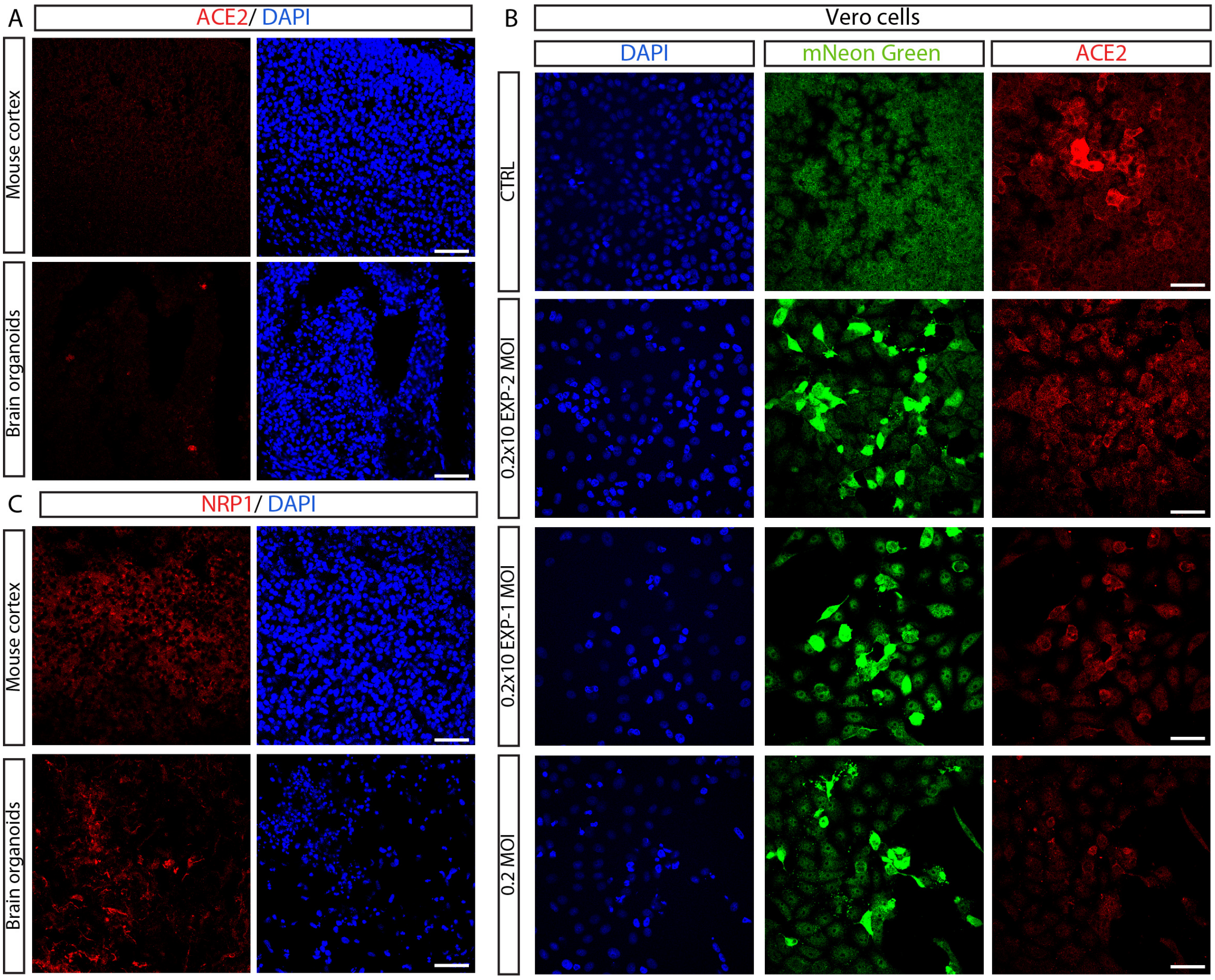


**Supplementary Figure S2. The ACE2 co-receptor NRP1 is expressed in human brain organoids *in vitro*.**

(A) Confocal images of mouse cortex (top) and brain organoids BOs (bottom) immunostained for ACE2 (in red). (B) Confocal images of CTRL and infected Vero cells with SARS-CoV2-mNeon Green modified virus (MOI: $0.5 \times 10^{-2}$; $0.5 \times 10^{-1}$; 0.5), immunostained with mNeon Green (in green) and ACE2 (in red). (C) Confocal images of mouse cortex (top) and brain organoids BOs (bottom) immunostained for NRP1 (in red). Counterstaining was performed using DAPI. Scale bars: 50 μm

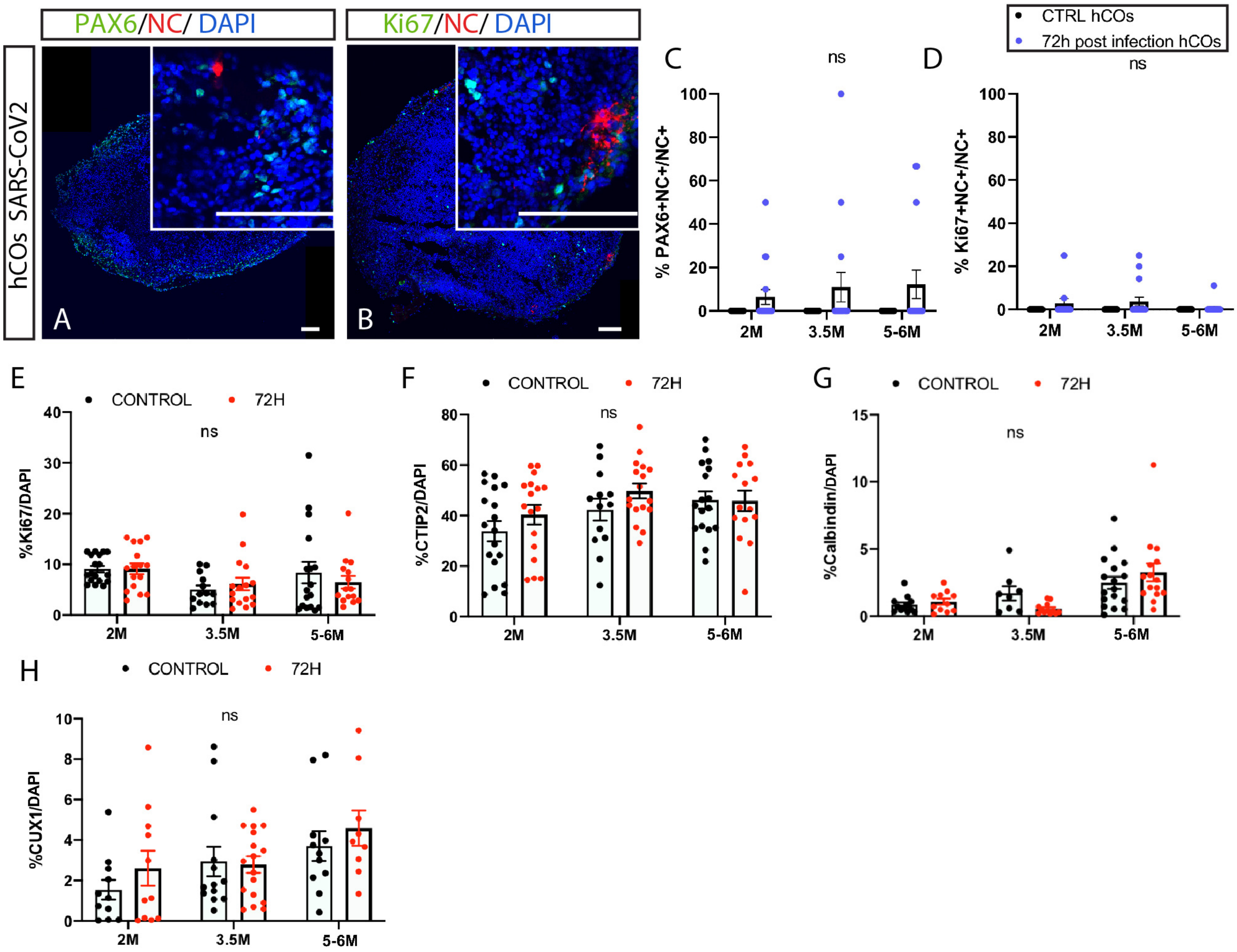


**Supplementary Figure S3. Progenitor populations are mostly non-infected and no effects on cell-type number are found following SARS-CoV2 infection.**

(A-B) Tile-Scan confocal images of cryosections immunostained for the SARS-CoV2 antigen nucleocapside (NC, in red) and either for the progenitor marker PAX6 (in green) or the proliferative marker Ki67 (in green) in 6M 72h post-infection hCOs. (C-D) Quantification of the percentage of double positive PAX6+NC+ (C); KI67+NC+ (D), among the total NC+ population in 2M, 3.5M and 5-6M 72h SARS-CoV2 infected hCOs compared to control (CTRL). (E-H) Quantification of the percentage of KI67+ (E), CTIP2+ (F), Calbindin+ (G), CUX1+ (H) cells among the DAPI+ population 72h post infection hCOs compared to CTRL hCOs. Data are represented as mean percentages ± SEM (2-3 differentiations: 2M n=11-18, 3.5M n=8-16, 5-6M n=17-11). Mixed-effect analysis with a Sidak's multiple comparison Ns= non-significant. Scale bars: 100µm (A-B).

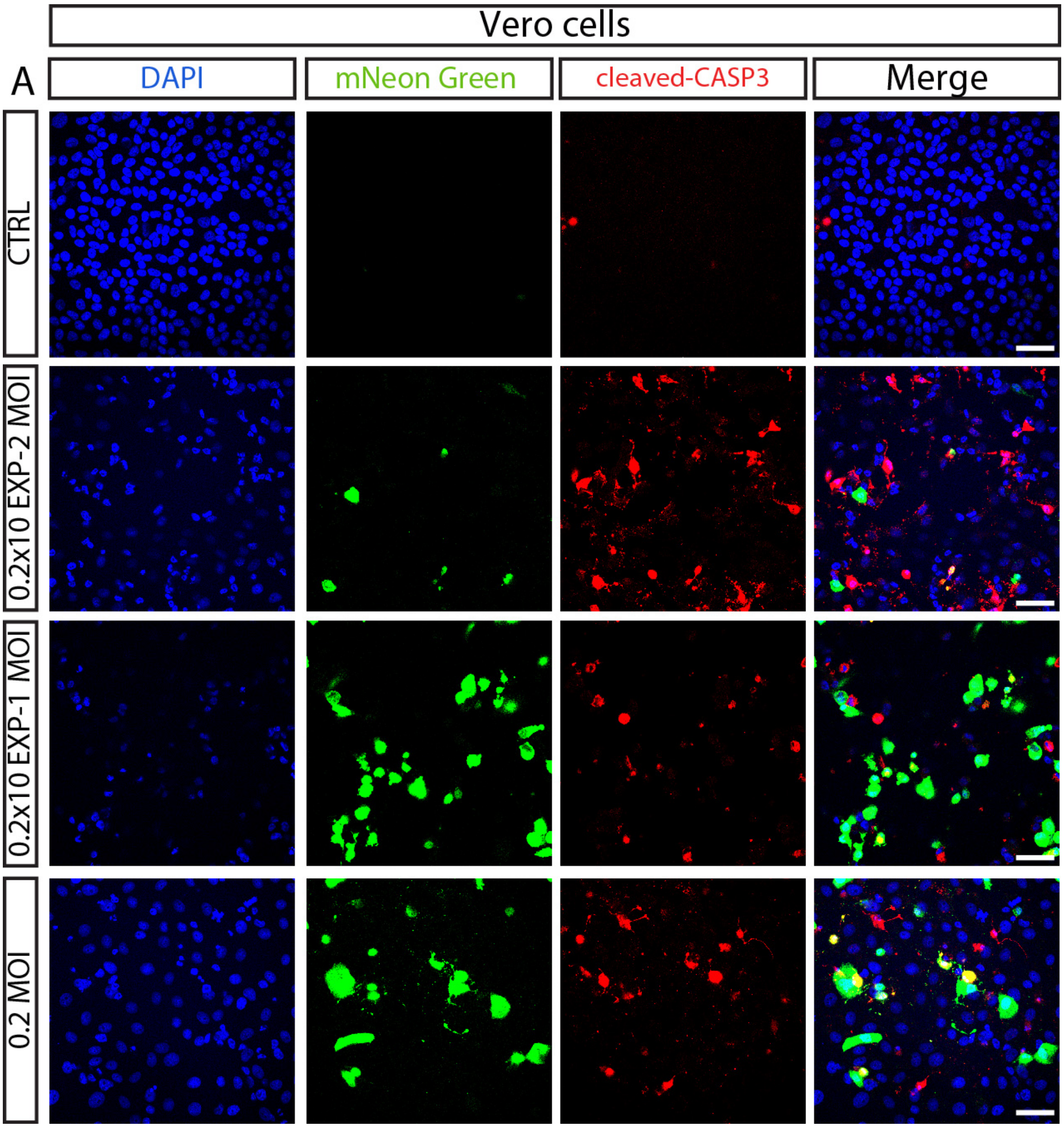


**Supplementary Figure S4. Vero cells show upregulation of the apoptotic cleaved-CASP3 marker following infectivity by SARS-CoV2.**

Confocal images of CTRL and infected Vero cells with SARS-CoV2-mNeon Green modified virus (MOI: $0.5x10^{-2}$; $0.5x10^{-1}$; 0.5), immunostained with mNeon Green (in green) and cleaved-CASP3 (in red). Counterstaining was performed using DAPI. Scale bars: 50 µm

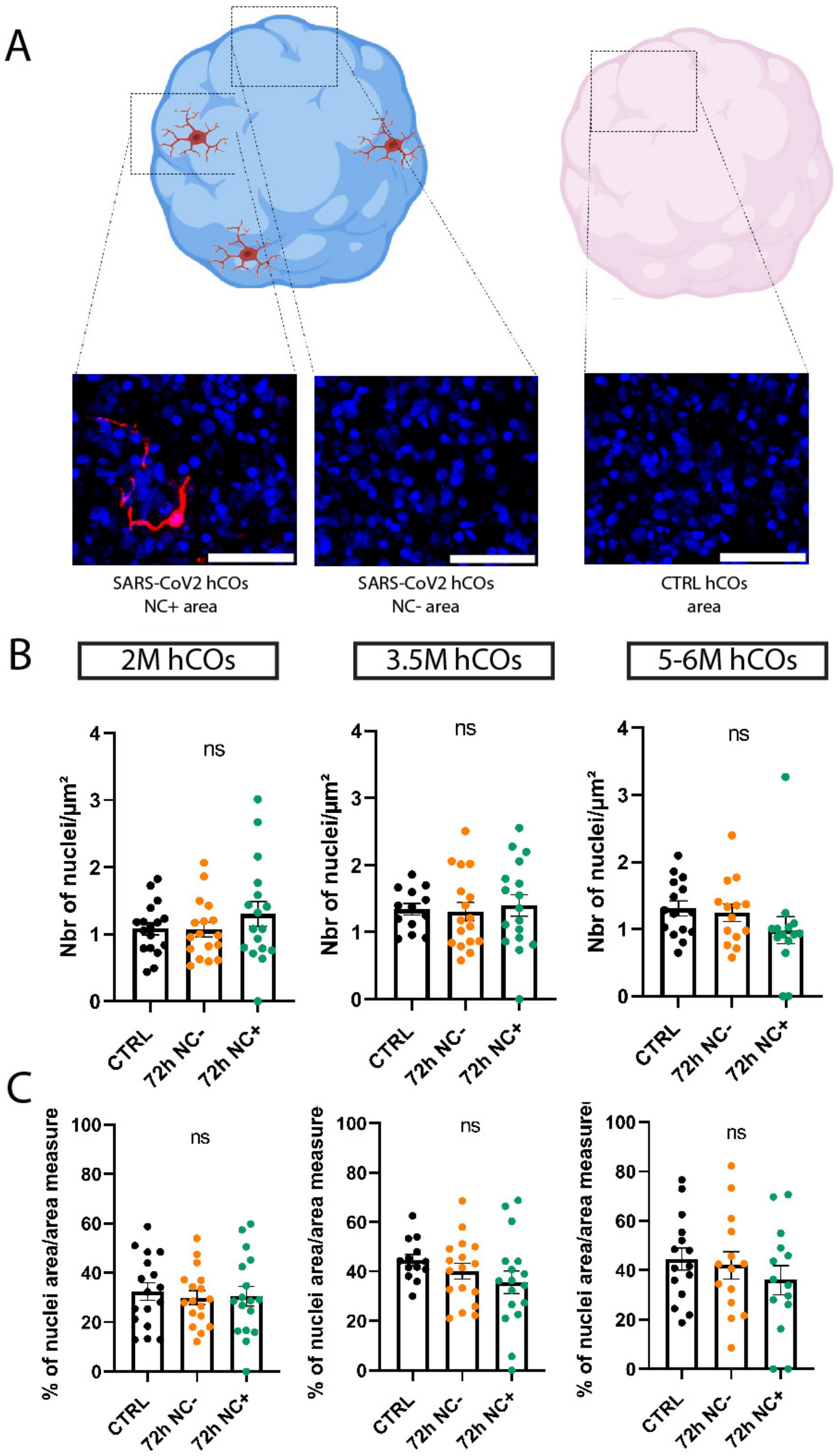

A
SARS-CoV2 hCOs
NC+ area
SARS-CoV2 hCOs
NC- area
CTRL hCOs
area
B
2M hCOs
3.5M hCOs
5-6M hCOs
Nbr of nuclei/µm²
ns
CTRL
72h NC-
72h NC+
C
% of nuclei area/area measure

**Supplementary Figure S5. Absence of cell death phenotypical-related changes in nuclei from NC+ infected areas in SARS-CoV2 hCOs**.
(A) Cartoon depicting the areas chosen for quantification of NC+ areas in 72h post infected hCOs (SARS-CoV2 COs NC+ area), in NC- areas in 72h post infected hCOs (SARS-CoV2 COs NC- area) and in CTRL hCOs (CTRL COs area). Created with BioRender.com. (B) Quantification of the nuclei DAPI+ cell density inside these areas at 2M, 3.5M and 5-6M hCOs. Data are represented as mean percentages ± SEM (3 differentiations: 2M n=17; 3.5M n=13-17 and 5-6M n=14-15). One-way ANOVA test with Tukey multiple comparison test (2M). Kruskal-Wallis test with Dunn multiple comparison test (3.5M and 5-6M). (C) Quantification of the (DAPI+) total nuclei area inside the areas measured at 2M, 3.5M and 5-6M hCOs. Data are represented as mean percentages ± SEM ( differentiations: 2M n=17; 3.5M n=13-17 at 3.5M 5-6M n=14-15). One-way ANOVA test with Tukey multiple comparison test (2M; 5-6M). Kruskal-Wallis test with Dunn multiple comparison test (3.5M) area. ns: non-significant. Scale bars: 50µm (A).

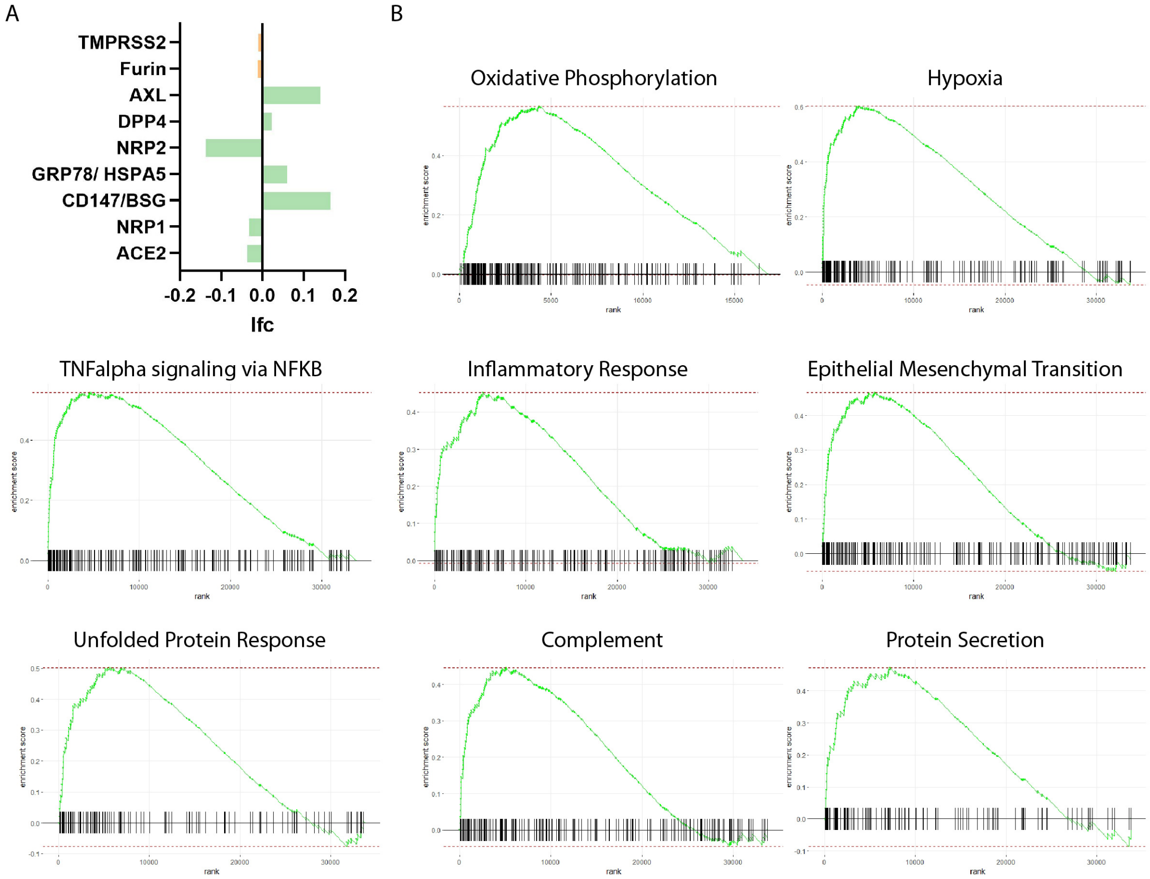


**Supplementary Figure S6. Cytokine release and transcriptomic profile of 72h post infected hCOs reveals upregulation of pro-inflammatory and cell survival genes.**

(A) No significant variation in the expression of receptors or co-receptors (in green) nor proteases (in orange) is observed comparing SARS-CoV2 to CTRL hCOs at 6M. (B) GSEA enrichment score curves for pathways involved in immune function and cell survival in SARS-Cov2 hCOs compared to CTRL hCOs. The y-axis represent the enrichment score and the x-axis shows the ranked list of genes, and the vertical bars on the x-axis show the genes that belong to gene set (title of each graph). The green line represent the genes enriched in the top ranking genes (enrichment profile). The red dotted line corresponds to the enrichment score (ES).

**Table S1. List of differentially expressed genes (DEG) in SARS-CoV2 infected COs compared to Control COs (padj<0.05)**

| ID | Gene | baseMean | Log2Foldchange | lfcSE | padj |
|---|---|---|---|---|---|
| ENSG00000196136 | SERPINA3 | 840,2491633 | 0,956961833 | 0,121082579 | 3,90E-11 |
| ENSG00000278233 | RNA5-8SN2 | 116,581409 | 0,017296461 | 0,019231072 | 3,11E-09 |
| ENSG00000278189 | RNA5-8SN1 | 39,71842579 | -0,017263545 | 0,019213535 | 5,18E-07 |
| ENSG00000205236 | AC105052.1 | 77,52020569 | -0,017263545 | 0,122436353 | 3,39E-05 |
| ENSG00000146648 | EGFR | 5418,080777 | -0,441833832 | 0,077785407 | 4,40E-05 |
| ENSG00000161714 | PLCD3 | 3213,095248 | 0,458844293 | 0,086407048 | 0,00030519 |
| ENSG00000197747 | S100A10 | 702,8735676 | 0,535202358 | 0,108587265 | 0,001785192 |
| ENSG00000205403 | CFI | 564,7945784 | 0,564086198 | 0,11901274 | 0,004281143 |
| ENSG00000205403 | IL13RA1 | 619,8942517 | 0,504626852 | 0,108860788 | 0,006820489 |
| ENSG00000112096 | SOD2 | 14040,64695 | 0,456754608 | 0,099675711 | 0,006843772 |
| ENSG00000167131 | CCDC103 | 26,31831263 | -0,310284001 | 0,079992506 | 0,006843772 |
| ENSG00000164929 | BAALC | 5045,678148 | 0,452584036 | 0,102333725 | 0,013079167 |
| ENSG00000242220 | TCP10L | 71,2160528 | 0,436310053 | 0,107416739 | 0,013079167 |
| ENSG00000273032 | DGCR5 | 689,155986 | -0,454460225 | 0,103458901 | 0,013079167 |
| ENSG00000026508 | CD44 | 14035,97654 | 0,444767837 | 0,102273512 | 0,01546958 |

| | | | | | |
|---|---|---|---|---|---|
| ENSG00000019991 | HGF | 714,4542456 | 0,497378612 | 0,115750446 | 0,01638698 |
| ENSG00000269897 | COMMD3-BMI1 | 169,897909 | -0,522425243 | 0,123405063 | 0,022567765 |
| ENSG00000187091 | PLCD1 | 3518,218292 | 0,365312297 | 0,08895687 | 0,037695238 |
| ENSG00000261408 | TEN1-CDK3 | 40,43499436 | -0,3432206 | 0,095188035 | 0,039358216 |

**Table S2. List of candidate pathways from enrichR represented from differentially expressed genes between CTRL and SARS-CoV2 infected COs (MSigDB Hallmark 2020)**

| Term | p-value | Adj p-value | Genes |
|---|---|---|---|
| Apoptosis | 4,51E-04 | 0,008905787 | HGF;SOD2;CD44 |
| Glycolysis | 8,48E-04 | 0,008905787 | EGFR;CD44;IL13RA1 |
| IL-6/JAK/STAT3 signaling | 0,003048397 | 0,021338777 | CD44;IL13RA1 |
| TNF-alpha signaling via NFKB | 0,015214448 | 0,063900683 | SOD2;CD44 |
| Estrogen response late | 0,015214448 | 0,063900683 | SERPINA3;CD44 |
| Reactive oxygen species pathway | 0,04555751 | 0,159451286 | SOD2 |
| Protein secretion | 0,087402649 | 0,173902367 | EGFR |
| Peroxisome | 0,094349807 | 0,173902367 | SOD2 |
| PI3K/AKT/mTOR signaling | 0,09521467 | 0,173902367 | EGFR |
| Coagulation | 0,123320172 | 0,173902367 | CFI |
| Fatty Acid Metabolism | 0,139949124 | 0,173902367 | S100A10 |
| UV Response up | 0,139949124 | 0,173902367 | SOD2 |
| Il2/STAT5 signaling | 0,173108927 | 0,173902367 | CD44 |
| Hypoxia | 0,173902367 | 0,173902367 | EGFR |
| P53 pathway | 0,173902367 | 0,173902367 | S100A10 |
| Estrogen response early | 0,173902367 | 0,173902367 | CD44 |
| Allograft rejection | 0,173902367 | 0,173902367 | EGFR |
| KRAS signaling up | 0,173902367 | 0,173902367 | SERPINA3 |
| Interferon gamma response | 0,173902367 | 0,173902367 | SOD2 |
| Apical junction | 0,173902367 | 0,173902367 | EGFR |
| Epithelial mesenchymal Transition | 0,173902367 | 0,173902367 | CD44 |

**Table S3. List of Gene Set Enrichment Analysis (GSEA) in SARS-CoV2 infected COs compared to Control COs.**

| Name | NES | padj | Nbr Genes/pathway |
|---|---|---|---|
| Oxidative phosphorylation | 2,923296 | 5,47E-26 | 200/200 |
| Hypoxia | 2,782357 | 8,41E-23 | 196/200 |
| mTORC1 signaling | 2,647516 | 1,26E-18 | 200/200 |
| TNFA signaling via NFKB | 2,563416 | 1,9E-17 | 190/200 |
| Glycolysis | 2,324447 | 7,17E-13 | 198/200 |
| MYC targetsV1 | 2,246841 | 8,98E-12 | 199/200 |
| Apoptosis | 2,243768 | 1,32E-09 | 156/161 |
| Coagulation | 2,233589 | 5,46E-09 | 119/138 |
| Unfolded protein response | 2,168832 | 3,96E-08 | 111/113 |
| Adipogenesis | 2,16512 | 2,77E-10 | 198/200 |
| P53 pathway | 2,159195 | 5,49E-10 | 197/200 |
| Epithelial mesechymal transition | 2,153913 | 6,25E-10 | 197/200 |
| Interferon gamma response | 2,103518 | 2,3E-09 | 192/200 |
| Cholesterol homeostasis | 2,095274 | 3,65E-06 | 74/74 |
| Myogenesis | 2,068647 | 1,43E-08 | 184/200 |
| Inflammatory response | 2,06364 | 1,77E-08 | 175/200 |
| Complement | 2,042529 | 5,63E-08 | 178/200 |
| Estrogen response late | 1,959863 | 1,23E-07 | 192/200 |
| Fatty acid metabolism | 1,871923 | 4,29E-06 | 154/158 |
| Mitotic spindle | -2,04437 | 1,55E-08 | 199/199 |
| E2F targets | -2,26806 | 1,21E-11 | 200/200 |
| G2M checkpoint | -2,38767 | 1,08E-13 | 200/200 |

**Supplemental Experimental Procedures**

**Human Pluritpotent stem cell culture**

Human ESC H9 line (WiCell) was cultured using standard procedures for feeder-free cultures (WiCell). Briefly, hESC cultures were maintained in mTeSR plus medium (StemCell Technologies ref: 100-0276) on geltrex (Thermo Fisher ref: A1413302)-coated plates and passaged twice a week in a ratio 1:3-1:6 using EDTA (Sigma ref:03690). hESC-H9 cultures were amplified about 1-3 weeks before starting each experiment (2-6 passages) from cryopreserved stocks (passages ~30-40) and cells were kept for a maximum of 10-15 passages before thawing a new vial. The H9 line was authenticated by WiCell upon acquisition. hESC cultures were routinely assayed for the expression of pluripotency markers by qPCR and immunofluorescence experiments, as previously described[1]. Mycoplasm screenings were performed every 3 months. H9 cultures were kept ruinously without antibiotics under good laboratory practices to ensure the absence of bacteria and or fungi contaminations in the cultures. Genomic characterization was performed by copy-number-assay (qPCR CNV) on genomic DNA (Thermo Fisher) from hESC-H9 cells amplified from cryopreserved stocks (passages ~ 30-40).

**Human COs differentiation**

Cortical brain organoids are developed following a modified version of the Sasai protocol[2]. hESC-H9 cell line (Wicell) was dissociated using accutase (Stem Cell Technologies ref: 7922) to obtain a suspension of individual cells that were seed in 96 well U bottom plates (Nunclon sphera ref: 15396123) 9000 cells per well. Plates were centrifuged 2min at 550 rpm to cluster the cells at the bottom of the well. Cells were cultured in DMEM-F12 (ThermoFisher ref: 31331-093) with 20% KO serum (Fisher ref: 10828028), supplemented with 1/100 non-essential amino acid (NEAA; ThermoFisher ref: 11140-035), 1/100 penicillin/streptavidin (Pen/strep; ThermoFisher ref: 15140122) and 100µm β-mercaptoethanol (Sigma ref: 444203) for 14 days. Morphogens are added to the medium for 14 days: 5µM SB (SB431542; Sigma ref: S4317) and 3µM IWR1 (Sigma ref: 681669). Rock inhibitor (Y-27632 2HC; Bio-Connect ref: S1049) is added on day 0 when seeding the cells at 20µM and then at 10µM from day 2 to day 6. From day 14 to day 28, medium is composed of DMEM-F12, 1/100 sodium pyruvate (Na Pyr; ThermoFisher ref: 11360-039), 1/100 NEAA, 1/100 N2 (ThermoFisher ref: 17502048), 1/100 Pen/strep, 1/150 BSA (BSA fraction V; ThermoFisher ref: 9048-46-8), 1/50 B27 without vitamin A (ThermoFisher ref: 11500446) and 100µm β-mercaptoethanol. Organoids were transferred to 6cm dishes (Greiner ref: CLS430196) when their area was around 0,7mm$^2$ and placed on orbital shaker (75 rpm). From day 28 to day 35, medium is composed of half DMEM-F12 and half Neurobasal (ThermoFisher ref: 21103-049) and supplemented with 1/200 Na Pyr, 1/200 NEAA, 1/200 N2, 1/200 Glutamax (ThermoFisher ref: 35050-038), 1/300 BSA, 1/100 Pen/strep, 1/50 B27 without vitamin A and 50µm β-mercaptoethanol. From day 35 till the end of the culture, the medium is similar to day 28 to d35 but supplemented with B27 with vitamin A (ThermoFisher ref: 11530536). Organoids were transferred to bioreactor tubes (Bioké ref: 2800005) when their area was around 1,4mm$^2$ and kept in culture in a bioreactor at 75 rpm (Bioké ref: 2800000). At day 70, 1% of matrigel (Corning ref: 354230) was added to the culture. Medium was changed every 2 or 3 days and pictures were acquired on the same days to assess the area and overall quality of the organoids till their transfer to the bioreactors.

**MGE brain organoids differentiation**

hMGE brain organoids were differentiated from hESC-H9 cell line (Wicell) following protocols previously published[3,4]. Briefly, accutase-dissociated cells were plated in a U-bottom ultra-low-attachment 96-well plate (Nunclon sphera ref: 15396123) 9000 cells per well in DMEM-F12 media, supplemented with 20% of knockout serum, 1% NEAA, 1% Glutamax and 50 μM of Rock Inhibitor. The plate was then centrifuged at 100g for 3 min and incubated overnight at 37 °C with 5% CO2. The next day (day 1), the media was complemented with 100 nM LDN-193189 (Bio-techne ref: 6053), 10 μM SB431542, and 2 μM XAV939 (StemCell ref: 72672). Organoids were cultured in this media from days 1 to 8. On day 8, organoids were transferred to ventral patterning media containing DMEM-F12, 1% N2 supplement, 1% NEAA, 1% Glutamax and 0,1% heparin (Sigma ref: H3393-25KU), 100nM SAG (EMD Millipore, ref: 566660) and 2.5μM IWP2 (Sigma, ref: I0536). Media was changed every other day until day 20. After 20 days of stationary culture, organoids in 96-well plate were transferred to ultra-low-attachment 6-well plate in neural differentiation media without vitamin A (NDM-A) and placed on a shaker at 75rpm. NDM-A media contained 1:1 mixture of DMEM-F12 media and Neurobasal media, supplemented with 0.5% N2 supplement, 1% B27 supplement without vitamin A, 1% Glutamax, 0.5% NEAA, 0.025% human insulin solution (Sigma ref: I9278-5ML), 50 μM β-mercaptoethanol, and 1% (v/v) Pen/Strep. The media was changed every 2 days.

**Thalamic organoid differentiation**

Thalamic brain organoids are developed following the Park protocol[5]. Briefly, hESC-H9 cell line (Wicell) was dissociated using accutase (Stem Cell Technologies ref: 7922) to obtain a suspension of individual cells that were seed in 96 well U bottom plates (Nunclon sphera ref: 15396123) 9000 cells per well. Plates were centrifuged 3min at 500 rpm to cluster the cells at the bottom of the well. Cells were cultured in DMEM-F12 (ThermoFisher ref: 31331-093) with 20% KO serum (Fisher ref: 10828028), supplemented with 1/100 non-essential amino acid (NEAA; ThermoFisher ref: 11140-035), 1/100 penicillin/streptavidin (Pen/strep; ThermoFisher ref: 15140122), 1/100 Glutamax (ThermoFisher ref: 35050-038) and β-mercaptoethanol (stock final 100μm; Sigma ref: 444203) for 16 days. Morphogens are added to the medium for 16 days, from day 0 to day 8: 100nM LDN (Bio-techne ref: 60553), 10μM SB (SB431542 Sigma ref: S4317), 4μg/ml insulin (Sigma ref: I9278-5ML). Then, from day 8 to day 16, different morphogens were added to the medium: 1M PD325901 (MedChem Express ref: HY-10254) and 30ng/ml BMP7 (Peprotech 120-03). Rock inhibitor (Y-27632 2HC; Bio-Connect ref: S1049) is added on day 0 when seeding the cells at 20μM and then at 10μM from day 2 to day 6. From day 16 to day 28, medium is composed of DMEM-F12, 1/100 sodium pyruvate (Na Pyr; ThermoFisher ref: 11360-039), 1/100 NEAA, 1/100 N2 (ThermoFisher ref: 17502048), 1/100 Pen/strep, 1/100 Glutamax, 1/150 BSA (BSA fraction V; ThermoFisher ref: 9048-46-8), 1/50 B27 without vitamin A (ThermoFisher ref: 11500446) and 100μm β-mercaptoethanol. On day 25, organoids were transferred to 6cm dishes and placed on orbital shakers (75 rpm). From day 28 to day 35, medium was composed of half DMEM-F12 and half Neurobasal (ThermoFisher ref: 21103-049) and supplemented with 1/200 Na Pyr, 1/200 NEAA, 1/200 N2, 1/200 Glutamax (ThermoFisher ref: 35050-038), 1/300 BSA, 1/100 Pen/strep, 1/50 B27 without vitamin A and 50µm β-mercaptoethanol. From day 35 till the end of the culture (day 60), medium was similar to day 28 to d35 but supplemented with B27 with vitamin A instead of without (Thermo Fisher ref: 11530536), 5µg/ml heparin (Sigma ref: H3393-25KU), 10% FBS (Fisher Scientific ref: 11573397) and 1% matrigel (Corning ref: 354230) were added to the medium. Medium was changed every 2 to 3 days.

**Cerebellum organoid differentiation**

Cerebellum brain organoids were developed following a modified version of the Sasai protocol[6]. hESC-H9 cell line (Wicell) was dissociated using accutase (Stem Cell Technologies ref: 7922) to obtain a suspension of individual cells that were seeded in 96 well U bottom plates (Nunclon sphera ref: 15396123): 9000 cells per well. Plates were centrifuged 1min at 500 rpm to cluster the cells at the bottom of the well. Cells were cultured in DMEM-F12 (ThermoFisher ref: 31331-093) with 20% KO serum (Fisher ref: 10828028), supplemented with 1/100 non-essential amino acid (NEAA; ThermoFisher ref: 11140-035), 1/100 penicillin/streptavidin (Pen/strep; ThermoFisher ref: 15140122), 1/100 Glutamax (ThermoFisher ref: 35050-038) and 100µm β-mercaptoethanol (Sigma ref: 444203) for 14 days. Morphogens were added to the medium for 14 days. From day 0: 10µM SB (SB431542; Sigma ref: S4317) and 7µg/ml insulin (Sigma ref: I9278-5ML), chemically defined lipid concentrate (1% Invitrogen). Then, from day 2, 50µg/ml FGF2 (Peprotech ref: 100-18b) was added. From day 7, the concentration of SB and FGF2 were reduced respectively to 5µM and 25µg/ml. Rock inhibitor (Y-27632 2HC; Bio-Connect ref: S1049) was provided on day 0 when seeding the cells at 20µM and then at 10µM from day 2 to day 6. From day 14 to day 28, medium was composed of DMEM-F12, 1/100 sodium pyruvate (Na Pyr; ThermoFisher ref: 11360-039), 1/100 NEAA, 1/100 N2 (ThermoFisher ref: 17502048), 1/100 Pen/strep, 1/100 Glutamax, 1/150 BSA (BSA fraction V; ThermoFisher ref: 9048-46-8), 1/50 B27 without vitamin A (ThermoFisher ref: 11500446) and 100µm β-mercaptoethanol. Insulin (7µg/ml) (Sigma ref: I9278-5ML) and 1% co-lipids were also added to the medium till day 21. On day 25, organoids were transferred in a 6cm dish and placed on orbital shaker (75 rpm). From day 28 to day 45, medium was composed of half DMEM-F12 and half Neurobasal (ThermoFisher ref: 21103-049) and supplemented with 1/200 Na Pyr, 1/200 NEAA, 1/200 N2, 1/200 Glutamax (ThermoFisher ref: 35050-038), 1/300 BSA, 1/100 Pen/strep, 1/50 B27 without vitamin A and 50µm β-mercaptoethanol. From day 45 till the end of the culture (day 60), culture medium was similar to day 28 to d45 but supplemented with 5µg/ml heparin (Sigma H3393-25KU), 10% FBS (Fisher Scientific 11573397) and 1% matrigel (Corning ref: 354230). Medium was changed every 2 to 3 days.

**VeroE6 cell culture**

VeroE6 cells (Epithelial cells from an African green monkey, ATCC # CRL-1586) were cultured in DMEM medium (Sigma ref: D5796) supplemented with 10% of fetal bovine serum (FBS, Gibco ref: A5256701), 2mM of L-glutamine, 100 units of penicillin and 100 g/l of streptomycin in a 75 cm culturing plate. Cells were passaged every 3 days with trypsin-EDTA 0.05% (Thermo Fisher Scientific ref: 25300054).

**SARS-CoV2 infection**

All experiments have been conducted by the GIGA viral platform (www.gigaviralvectors.uliege.be) in a Biosafety Level 3 lab using the SARS-CoV2 strain (BetaCov/Belgium/GHB-03021/2020 (EPI ISL 407976|2020-02-03)[7] wild or recombinant SARS-CoV-2 strain harboring a mNeon-Green reporter[8]. VeroE6 cells were used to generate the SARS-CoV2 viral stock. Plaque assay method was used to determine the virus titer or plaque-forming unit (PFU)[9].

For infection of VeroE6 cells, 50.000 cells were plated on 18 mm round glass coverslips (Carl Roth ref: P233.1) in 24 well-plates and cultured in DMEM with 10% FBS. Twenty-four hours after plating when VeroE6 cells were at about 80% confluency cells were infected with $10^2$; $10^3$; $10^4$ PFU of SARS-CoV2 virus (corresponding to an expected $0.2x10^{-2}$; $0.2x10^{-1}$; 0.2 MOI, respectively) for 2-3h at 37°C. After infection the virus was removed, cells were washed with PBS and medium was added until the indicated time post-infection. After 72h post infection, cells were washed with PBS x3 and either fixed with 4 % PFA for 30 min for immunostaining experiments or collected for total RNA extraction in RLT buffer (Qiagen) and extracts were frozen at -80°C until further use.

For infection of cortical, MGE, THL and CRB organoids, organoids were placed in 96 U bottom plates and incubated with SARS-CoV2 corresponding to an expected 0.5 MOI for 2-3h in 100µl of culture medium at 37°C. The number of cells inside the organoids was assessed by measuring the area of organoids from phase contrast images acquired from live organoids and extrapolating the corresponding number of cells according to previous scale/measurements performed in the lab from single cell dissociations of organoids. When indicated a MOI 1 was used to infect cortical organoids following incubation with SARS-CoV2 virus. After infection the virus was removed, organoids were washed with PBS and kept in 100µl of culture medium for 24h or 72h. After 24h or 72h post-infection organoids were either fixed with PFA 4% 1h at RT, washed 1-2 times with PBS and let in sucrose 30% for immunostaining, kept in PFA 4% for 24h for clarity experiments, or collected for total RNA extraction in RLT buffer (Qiagen) and extracts were frozen at -80°C until further use.

**Clear, unobstructed brain/body imaging cocktails and computational analysis (CUBIC) protocol**

Clarification of the whole organoid was performed based on the CUBIC method of Tainake[10]. Organoids were first fixed in PFA 4% overnight at 4°C. After fixation, they were washed in PBS 3 times 2h at RT with agitation and then immersed in half water and half CUBIC-L (10% N-butyldiethanolamine (Sigma ref: 471240) and 10% TritonX-100 (Sigma ref: 93443) for 6h at 37°C with agitation. Organoids were then immersed in CUBIC-L for 5 days at 37°C with agitation and CUBIC-L was refreshed every 2 days. Then 3 washes of 2h at RT with agitation were performed. Organoids were blocked (10% donkey serum (Bio-Connect ref: 017-000-121), 1% TritonX-100, 0,2% sodium azide (Sigma ref:

26628-22-8) in PBS) for 1 day at 4°C with agitation. Primary antibodies (nucleocapside 1/500; Sino Biol ref: 40143-MM08; GFP 1/1000; Abcam ref: ab6673) were added in antibodies solution (1% donkey serum, 0,2% TritonX-100, 0,2% sodium azide in PBS) for 2 days at 4°C with agitation. Organoid were washed 3 times 1h in wash solution (3% NaCl, 0,2%Tritonx-100 in PBS) at RT and kept overnight at 4°C before the addition of secondary antibodies. Incubation with anti-mouse 555 (ThermoFisher ref: A-31570) and anti-goat 488 (ThermoFisher ref: A-11055) was done in antibodies solution for 2 days at 4°C with agitation. The wash step in wash solution was repeated and organoids were then washed in PBS 3 times 15min with agitation at RT. To clarify the organoids, they were put in half water and half CUBIC-R (water 25 ml, antipyrine 39,3 gr, Nicotinamide 26,25 gr. RI=1.5444) for 1 day at 37°C with agitation, then, immersed in 100% CUBIC-R for 2 days at 37°C with agitation. Imaging was done the day after using a Zeiss Lightsheet Z1 microscope. Images were processed with the Imaris software (Oxford Instruments) using a spot analysis with a manual threshold. Movies were also generated via the function movies of Imaris.

**Cryosectioning of human cortical organoids**

Organoids were rinsed in phosphate-buffered solution (PBS), fixed in 4% paraformaldehyde PBS [PAF] for 45 minutes at 4 °C, then transferred to 30% sucrose solution in PBS for 48h. Excessive sucrose solution was removed from organoids by moving them through embedding medium (Epredia, Richard-Allan Scientific, MI, USA) after which they were flash frozen in blocks of embedding medium on dry ice and stored at -80°C until further use. Blocks were cut on a Cryostar NX70 cryostat into 14μm thick sections collected onto slides (Superfrost Plus Adhesion, Epredia Ref. J1800AMNZ), 1 to 4 series per slide, air-dried overnight, and kept at −20°C until staining.

**Immunofluorescence and Confocal Imaging**

Immunofluorescence of organoid cryosections was performed directly on the slides as described before[11]. For all stainings slides were first rinsed in PBS for 5 min to remove embedding medium followed by antigen retrieval in target retrieval solution (Dako ref: S1699) at 95 °C for 15 minutes. Slides were then rinsed 3 x 5min in a bath of PBS at room temperature. Permeabilization was done in 5% BSA (Carl Roth ref: 0163.2), 5% donkey serum, and 0.3% Triton X-100 in PBS for 30 min at room temperature by adding 500 μL of solution to each slide. Afterwards, primary antibodies were added in a buffer solution of 3% BSA, 1% donkey serum, and 0.1% Triton, in PBS, at 350 μL per slide and slides were incubated at 4°C overnight in a wet box and covered with parafilm. The next day, slides were washed 3× in PBS and incubated for 3h in a dark box at RT with secondary antibodies in buffer solution at 1:500. Slides were then washed 2x 5min with 500 μL PBST (0.1% Tween-20 (Sigma ref: 9036-19-5) in PBS) and 1x with PBS and then incubated in DAPI solution (2 ng/ml) for 20 min. Slides were washed 2x in PBS, cover-slipped using fluorescent mounting agent (Dako ref: S3023, Agilent Technologies Inc, Santa Clara, CA, USA), and air-dried for 48h. Immunostained slides were stored at 4°C for imaging and 20°C for long-term storage. All primary and secondary antibodies used are listed in the tables below.

Immunostained organoid sections were imaged with a 20X, 40X and 60X plan-apochromat objective on a Nikon A1R confocal microscope. XY scanning stage was used for tile scans with 405 nm, 488 nm, 555 nm, and 647 nm lasers.

| Primary antibodies | Species | Company | Catalog number | Dilution |
|---|---|---|---|---|
| CALB | Rabbit | Swant | CB-38a | 1/1000 |
| CASPASE3 | Rabbit | Cell Signaling | 9661 | 1/400 |
| CTIP2 | Rat | Abcam | ab18465 | 1/300 |
| CUX1 | Rabbit | Milipore | ABE217 | 1/150 |
| GFAP | Rabbit | Dako | z0334 | 1/500 |
| H2AXγ | Rabbit | Cell Signaling | 2577S | 1/500 |
| KI67 | Rabbit | Abcam | ab15580 | 1/500 |
| MAP2 | Chicken | Abcam | ab5392 | 1/2000 |
| NRP1 | Rabbit | Abcam | ab81321 | 1/500 |
| NUCLEOCAPSIDE SARS-COV2 | Mouse | Sino Biological | 40143-MM08 | 1/500 |
| PAX6 | Rabbit | Covance | PRB-278P | 1/500 |
| SERPINA3 | Rabbit | Sigma | HPA002560 | 1/1000 |
| SOD2 | Mouse | Invitrogen | MA1106 | 2μg/ml |

| Secondary antibodies | Species | Company | Catalog number | Dilution |
|---|---|---|---|---|
| Anti chicken 647 | Donkey | Jackson ImmunoResearch | 703-605-155 | 1/500 |
| Anti mouse 555 | Donkey | Invitrogen | A31570 | 1/500 |
| Anti rabbit 488 | Donkey | Invitrogen | A21206 | 1/500 |
| Anti rabbit 647 | Donkey | Invitrogen | A31573 | 1/500 |
| Anti rat 488 | Donkey | Jackson ImmunoResearch | 712-545-150 | 1/500 |

**Quantification of the percentage of NC, KI67, PAX6, CTIP2, CUX1, CALB, GFAP, caspase3, SERPINA3, SOD2, H2AXγ**

Image quantifications were done in ImageJ/Fiji. NC, GFAP, SOD2 and caspase3 were counted manually due to the low number of cells positive for each staining. Ki67 and CTIP2 were counted automatically using an ImageJ/Fiji macro. The optimal threshold for each staining was assessed by trial error and was tested on different organoids and different stages (in control and infected organoids) for confirmation of the accuracy of the threshold used. Triangle threshold was used for KI67 with a particles size of 20 $\mu m^2$. For CTIP2, as the staining leads to different brightness levels of positive neurons, the picture was first smoothed 2 times, background was subtracted and the default threshold was used. A binary watershed was applied and particles were counted (size 2$\mu m^2$ onwards). The stainings PAX6, CUX1, CALB, SERPINA3, SOD2 and H2AXγ were counted semi-automatically with the threshold set manually. The threshold used for all quantifications was moments except for the quantification of CALB (default threshold). The choice of the threshold value was made in each case by comparing the generated binary picture (white and black) with the original picture (16 bits) and setting the value that would match best the signal observed. Particle size choice was 15$\mu m^2$ for CALB, 10$\mu m^2$ for PAX6, 20$\mu m^2$ for CUX1, 20$\mu m^2$ for SERPINA3 and 5$\mu m^2$ for H2AXγ.

**Quantification of the percentage of double positive NC/KI67, NC/PAX6, NC/CTIP2, NC/CUX1, NC/CALB, NC/GFAP, NC/caspase3, SOD2/GFAP cells**

Image quantifications were done using ImageJ/Fiji. Double positive cells for NC and another cell marker were assessed manually by checking the presence of the specific cell marker on top of the NC signal. Double positive cells for SOD2 and GFAP were also assessed manually with the help of the DAPI staining.

**Quantification of GFAP+ organoid area**

Quantification of GFAP+ organoid area was done automatically using a ImageJ/Fiji macro as performed before[11]. The first step of the macro is to define the whole area of the organoid. This step is done based on the DAPI signal using the Huang threshold to capture the maximal signal, to have the full organoid recognized as one object. The second step is to define a mask from this object and to apply it on the channel corresponding to the GFAP. The threshold used for the GFAP signal was moments. The third step is to measure the percentage of organoid stained by GFAP with the function "%Area fraction" which give this percentage of the organoid stained.

**Total cell density quantification**

Total cell density is expressed as the total number of DAPI+ cells divided by the organoid area. The number of DAPI cells was counted using an ImageJ/Fiji macro for which the parameters were tested on different organoids from different stages and from both conditions (control or infected). Otsu was the threshold used and the particle size used was 20$\mu m^2$. The area of organoids was also assessed via an ImageJ/Fiji macro where the DAPI signal is detected using the Huang threshold to capture the maximal signal, to recognize the full organoid as one object. Next, the area of the organoid was measured via the "Area" function.

**Localized nuclei density and nuclei area quantification**

Localized nuclei density and nuclei area were assessed manually in CTRL and 72h infected hCOs. Measurements in the SARS-CoV2 NC+ Area in 72h hCOs: An arbitrary square size was drawn on the top of an infected area in 72h hCOs, and the number of nuclei as well as their area were measured manually. Measurements in the SARS-CoV2 NC- Area in 72h hCOs: A similar square size was drawn outside of the infected area in the same infected organoid and number of nuclei and their area were measured manually. Measurements in the CTRL hCOs Area: A similar square size was drawn in a similar region as the one chosen in 72h infected hCOs and number of nuclei and their area were measured manually.

### RNA extraction, RNA sequencing, Data Analysis and Differential Expression Analysis

RNA was extracted using the RNeasy mini kit (Qiagen ref: 74104), followed by a DNAse treatment (Ambion ref: AM2222). Sample quality was assessed using a Nanodrop 2000 (Thermo Scientific). High quality RNA samples RIN values (8-10) were used for sequencing. 4 samples of CTRL hCOs from two independent differentiations and 4 samples from SARS-CoV2 72h infected hCOs from two different experiments were analyzed. The total extracted RNA was analyzed by the GIGA Genomic platform (https://www.gigagenomics.uliege.be) and generated the polyA RNA-seq libraries using Illumina Stranded Total RNA Prep Ligation with Ribo-Zero Plus following the manufacturer protocol (Illumina ref: 20040529). The libraries were dosed through qPCR using the KAPA SYBR Fast* Mastermix (2X) kit Universal (Sopachem ref: KK4602) and were sequenced using the Illumina Novaseq 6000 sequencer with the flowcell Novaseq S4 V1.5 (300 cycles), generating pair end 2x150bp reads. Raw reads were processed through nf-core RNAseq pipeline (version 3.0) using human genome build GRCh38 and the annotations from Ensembl release 103 (ensemble.org).
For the analysis of the expression of SARS-CoV2-related genes, raw reads were mapped to the virus reference sequence (NCBI Reference Sequence: NC_045512.2).
Downstream analysis was performed on R using DESeq2 method for normalization, clustering and differential expression analysis.

### Bioinformatic analysis

Volcano plot was generated on R Studio from the DEG files and pajd value threshold for significance was set at 0.05. Gene Set Enrichment Analysis (GSEA) analysis was also performed on R Studio using the fsgea package[12]. Pathways used in the analysis are the one from Human MSigDB Collections (H: Hallmark gene sets) downloaded from https://www.gsea-msigdb.org/gsea/msigdb/human/collections.jsp. Threshold was set up at 0.00001 from the padj value to take into account the enriched pathways. GSEA was also perform with the desktop GSEA application (https://www.gsea-msigdb.org/gsea/index.jsp) and give similar results to fgsea on R Studio.

### Statistical analysis

All statistical analysis was performed using GraphPad (Prism). All data were first tested for normality with a Shapiro-Wilcoxon test and based on the result of this test, the parametric or corresponding non-parametric version of the test was used. The test for multiple comparisons was chosen based on the recommendation from GraphPad Prism. All information concerning the test used and the multiple comparisons test applied are found in the legend of the corresponding figure. Normalization of the data has been done by trial, within each trial, each individual value is divided by the mean of all sample values of each trial. All data presented are from at least 3 independent batches. N.s.= non-significant; * p<0.05; ** p<0.01; *** p<0.001; **** p<0.0001
